# Two unique biological response-modifier glucans beneficially regulating gut microbiota and faecal metabolome in a non-alcoholic steatohepatitis animal model, with potential for applications in human health and disease

**DOI:** 10.1101/2022.06.23.497433

**Authors:** Senthilkumar Preethy, Nobunao Ikewaki, Gary A Levy, Kadalraja Raghavan, Vidyasagar Devaprasad Dedeepiya, Naoki Yamamoto, Subramaniam Srinivasan, Natarajan Ranganathan, Masaru Iwasaki, Rajappa Senthilkumar, Samuel JK Abraham

## Abstract

**Objective:** The gut microbiome and its metabolites, influenced by age and stress, reflect the metabolism and immune system’s health. We assessed the gut microbiota and faecal metabolome in a Stelic Animal Model of non-alcoholic steatohepatitis (NASH).

**Design:** This model was subjected to the following treatments: reverse osmosis water, AFO-202, N-163, AFO-202+N-163, and telmisartan. Faecal samples were collected at 6 weeks and 9 weeks of age. The gut microbiome was analysed using 16S ribosomal RNA sequence acquired by next-generation sequencing and the faecal metabolome using gas chromatography-mass spectrometry.

**Results:** The gut microbial diversity increased greatly in the AFO-202+N-163 group. Post-intervention, the abundance of Firmicutes decreased, while that of Bacteroides increased and was the highest in the AFO-202+N-163 group. The decrease in the Enterobacteria and other Firmicutes abundance and in the Turicibacter and Bilophila abundance was the highest in the AFO-202 and N-163 groups, respectively. The Lactobacillus abundance increased the most in the AFO-202+N-163 group. The faecal metabolites spermidine and tryptophan, beneficial against inflammation and NASH, respectively, were greatly increased in the N-163 group. Succinic acid, beneficial in neurodevelopmental and neurodegenerative diseases, increased in the AFO-202 group. Decrease in fructose was the highest in the AFO-202 group. Leucine and phenylalanine decreased, whereas ornithine, which is beneficial against chronic immune-metabolic-inflammatory pathologies, increased in the AFO-202+N-163 group.

**Conclusion:** AFO-202 treatment in mice is beneficial against neurodevelopmental and neurodegenerative diseases and has prophylactic potential against metabolic conditions. N-163 treatment has anti-inflammatory effects against organ fibrosis and neuroinflammatory conditions. In combination, they present anticancer activity.

**Key messages:**

- The influence of gut microbiome on fecal metabolome and their association to several diseases is already known.
- This study proves the efficacy of 1,3-1,6 beta glucans with pre-biotic potentials, beneficially influencing both gut microbiome and metabolome.
- These results recommends for an in-depth exploration of relationship among pre-biotics, gut microbiome and gut-multi-organ axes on the fundamentals of disease onset.
- Hidden prophylactic and therapeutic solutions to non-contagious diseases with Aureobasidium pullulans produced 1,3-1,6 beta glucans may be unveiled.

## INTRODUCTION

With approximately 100 trillion microorganisms present in the human gastrointestinal tract, the microbiome is now considered a virtual organ of the body. The microbiome encodes over three million genes producing thousands of metabolites as compared to the 23,000 genes of the human genome, thus replacing several host functions and influencing the host’s fitness, phenotype, and health. The gut microbiota influences several aspects of human health, including the immune, metabolic, and neurobehavioural traits.[1] The gut microbiota ferments non-digestible substrates such as dietary fibres and endogenous intestinal mucus, which support the growth of specialist microbes producing short-chain fatty acids (SCFAs) and gases. The major SCFAs produced are acetate, propionate, and butyrate. Butyrate is essential for maintaining colonic cells, apoptosis of colonic cancer cells, activation of intestinal gluconeogenesis, maintenance of oxygen balance in the gut, and prevention of gut microbiota dysbiosis and has beneficial effects on glucose and energy homeostasis. Propionate is transported to the liver, where it regulates gluconeogenesis; acetate is an essential metabolite for the growth of other bacteria and regulates central appetite.[1] Gut dysbiosis, i.e. altered state of the microbiota community, is associated with several diseases, including diabetes, metabolic disorders, obesity, cancer, rheumatoid arthritis, and neurological disorders such as Parkinson disease (PD), Alzheimer disease, multiple sclerosis (MS), and autism spectrum disorders (ASD).[2, 3] The faecal metabolome represents the functional readout of the gut microbial activity; it can be considered an intermediate phenotype mediating the host– microbiome interactions. On an average, 67.7 ± 18.8% of the faecal metabolome variance represents the gut microbial composition. Thus, faecal metabolic profiling is a novel tool to explore the association between microbiome composition, host phenotypes, and disease states.[4] Other than faecal microbiota transplantation, probiotics and prebiotic nutritional supplements can help in restoring the dysbiotic gut to a healthy state.

Beta-glucans are one of the most promising nutritional supplements with established efficacy against metabolic diseases, diabetes, cancer, cardiovascular diseases, and neurological diseases. Beta-glucans derived from two strains of the black yeast Aureobasidium pullulans, AFO-202 and N-163, have beneficial effects against diabetes,[5] dyslipidaemia,[6] ASD,[7, 8] Duchenne muscular dystrophy,[9] non-alcoholic steatohepatitis (NASH),[10] and infectious diseases including coronavirus disease (COVID-19).[11, 12] A previous study showed that AFO-202 beta-1,3-1,6 glucan balanced the gut microbiome in ASD children.[13] In a study on the Stelic Animal Model (STAM) of NASH, AFO-202 beta-glucan significantly decreased the inflammation-associated hepatic cell ballooning and steatosis, whereas N-163 beta-glucan decreased fibrosis and inflammation. The combination of AFO-202 and N-163 significantly decreased the non-alcoholic fatty liver disease (NAFLD) activity score (NAS).[10] This study was undertaken as an extension of the NASH study to assess the faecal microbiome and metabolome profile before and after administration of AFO-202 and N-163 beta-glucans individually and in combination.

## METHODS

### Mice

This study was conducted in accordance with the Animal Research: Reporting of In Vivo Experiments guidelines. C57BL/6J mice were obtained from Japan SLC Inc. (Tokyo, Japan). The animal care followed the following guidelines: Act on Welfare and Management of Animals (Ministry of the Environment, Japan, Act No. 105; 1 October 1973), Standards relating to the care and management of laboratory animals and relief of pain (Notice No. 88 of the Ministry of the Environment, Japan; 28 April 2006), and Guidelines for proper conduct of animal experiments (Science Council of Japan; 1 June 2006). Protocol approval was obtained from the SMC Laboratories, the Japanese equivalent of the Institutional Animal Care and Use Committee (Study reference no.: SP_SLMN128-2107-6_1). The mice were maintained in a specific pathogen-free facility under controlled conditions: temperature, 23 ± 3°C; humidity, 50 ± 20%; 12-h artificial light and dark cycles (light from 8:00 to 20:00); and adequate air exchange.

The STAM of NASH was developed as previously described.[10] A single subcutaneous streptozotocin injection of 200 µg (STZ, Sigma-Aldrich, USA) was administered 2 days after birth. The mice were fed a high-fat diet (HFD, 57 kcal% fat, Cat# HFD32, CLEA Japan, Inc., Japan) from 4 to 9 weeks of age. All mice developed liver steatosis and diabetes, and at 3 weeks, steatohepatitis was observed histologically.

### Study groups

The mice were randomly allocated to the following five study groups (n = 8 in each):

**Vehicle group/control group:** The mice in this group were orally administered 5 mL/kg of reverse osmosis water as the vehicle solution once dailyfrom 6 to 9 weeks of age.
**AFO-202** beta-glucan group: The mice in this group were orally administered 1 mg/kg of AFO-202 beta-glucan supplemented in 5 mL/kg of the vehicle once daily from 6 to 9 weeks of age.
**N-163** beta-glucan group: The mice were orally administered 1 mg/kg of N-163 beta-glucan supplemented in 5 mL/kg of the vehicle once daily from 6 to 9 weeks of age.
**AFO-202+N-163** beta-glucan group: The mice were orally administered 1 mg/kg of AFO-202 supplemented in 5 mL/kg of the vehicle once daily as well as 1 mg/kg of N-163 supplemented in 5 mL/kg of the vehicle once daily from 6 to 9 weeks of age.
**Telmisartan** group: The mice were orally administered 10 mg/kg of telmisartan in the vehicle once daily from 6 to 9 weeks of age.

### Test substances

The AFO-202 and N-163 beta-glucans were provided by GN Corporation Co. Ltd., Japan. Telmisartan (Micardis) was purchased from Boehringer Ingelheim GmbH, Germany.

### Randomisation

The NASH model mice were randomised into the aforementioned groups (n = 8 each) at 6 weeks of age based on their body weight on the day before starting the treatment. Randomisation was performed by body weight-stratified random sampling using Excel software. They were stratified by body weight to obtain the standard deviation, and the difference in the mean weights among the groups was negligible.

### Animal monitoring and sacrifice

Viability, clinical signs (lethargy, twitching, and laboured breathing), and behaviour of the mice were monitored daily. Body weight was recorded daily before commencing the treatment. The mice were observed for significant clinical signs of toxicity, moribundity, and mortality before and after administration of the solutions. The animals were sacrificed at 9 weeks of age by exsanguination through direct cardiac puncture under isoflurane anaesthesia (Pfizer Inc., USA.).

### Collection of faecal pellets samples

Frequency: The faecal samples were collected at 6 weeks of age (before treatment administration) and 9 weeks of age (before sacrifice). Procedure: At 6 weeks of age, faecal samples were collected from each mouse using the clean catch method. The animals were handled using clean gloves sterilised with 70% ethanol. The abdomen was gently massaged, and the bottom of the mouse was positioned over a fresh sterilised petri dish to collect 1-2 faecal pellets. At the time of sacrifice, faecal samples were aseptically collected from the caecum. The tubes containing the faeces were immediately placed on ice. These tubes were snap-frozen in liquid nitrogen and stored at −80°C for shipping. Figure 1 shows the groups and corresponding faecal sample numbers allotted for microbiome and metabolome analyses.

**Figure 1:**
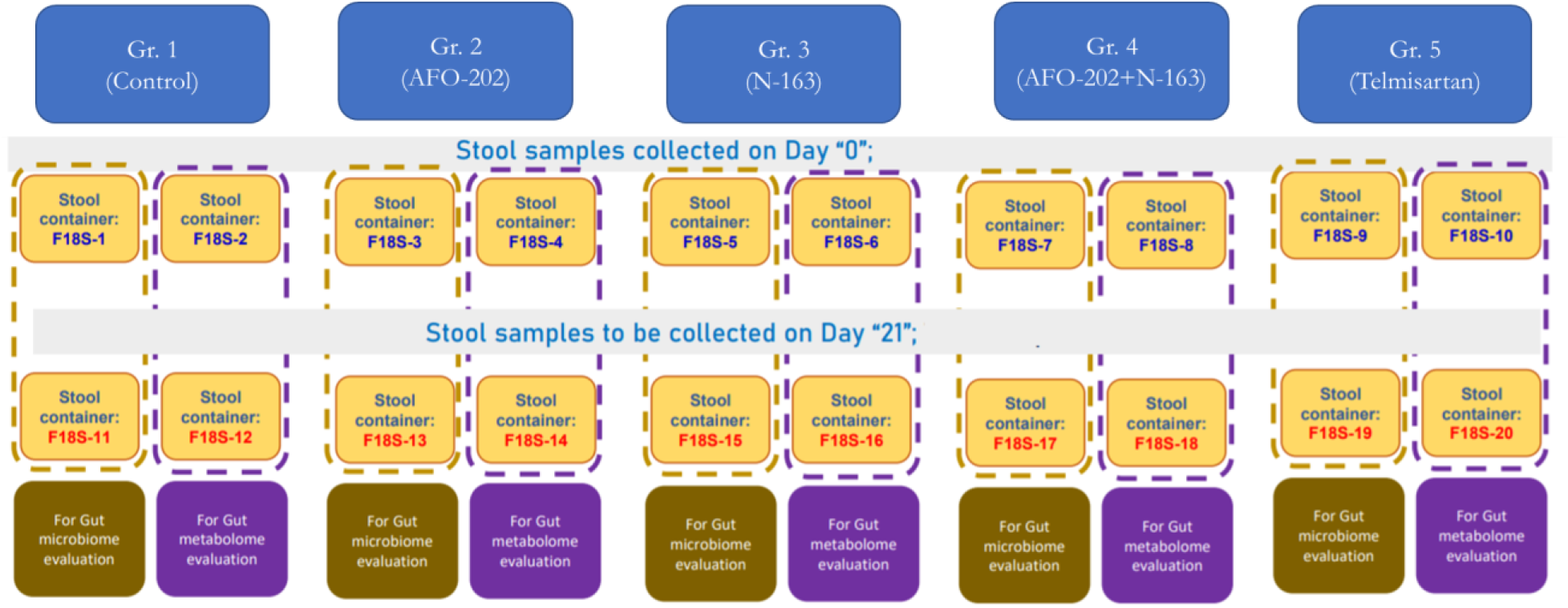
Study groups and faecal samples analysis reference numbers allotted for gut microbiome and metabolome analysis

### Microbiome analysis

In this analysis, the 16S ribosomal RNA (rRNA) sequence data acquired by next-generation sequencing (NGS) from faecal RNA were used to perform community analysis using Quantitative Insights Into Microbial Ecology (QIIME2) program for microbial community analysis. The raw read data in FASTQ format output from NGS were trimmed to remove adapter sequences and low-QV regions that may have been included in the data. Cutadapt was used to remove adapter sequences from DNA sequencing reads. Trimmomatic was used as a read-trimming tool for Illumina NGS data. The adapter sequence was trimmed using the adapter trimming program Cutadapt if the trimming of the region at the end of the read sequence overlapped the corresponding sequence by at least one base (mismatch tolerance: 20%). When reads containing N were present in at least one of Read1 and Read2, both Read1 and Read2 were removed.

Illumina adapter sequence information:

Read1 3’ end side

CTGTCTTCTATACACATCTCCGAGCCCACGAGAC

Read2 3’ end side

CTGTCTTCTATACACATCTGACGCTGCCGACGA

Trimming of the low-QV regions was performed on the read data after processing, using the QV trimming program Trimmomatic under the following conditions:

The window of 20 bases was slid from the 5’ side, and the area with average QV <20 was trimmed.

After trimming, only the reads with >50 bases remaining in both Read1 and Read2 were used as outputs.

### Population analysis

The microbial community analysis based on the 16S rRNA sequence was performed on the sequence data trimmed in the previous section using QIIME2. The annotation program “sklearn” of QIIME2 was used to annotate the amplicon sequence variant (operational taxonomic units) (ASV [OTU]) sequences.

Using “sklearn,” the ASV (OTU) sequences obtained were annotated with taxonomy information, i.e. Kingdom/Phylum/Class/Order/Family/Genus/Species, based on the 16S rDNA database.

The dataset of the 16S rDNA database “greengenes” provided on the QIIME2 resources site was used for the analysis. The ASVs (OTUs) obtained were aggregated and graphed based on the taxonomy information and the read counts of each specimen. Based on the composition of the bacterial flora for each specimen compiled above, various index values for alpha-diversity were calculated.

### Metabolome analysis

After lyophilisation of the faecal sample, approximately 10 mg of the sample was separated and extracted using the Bligh-Dyer method, and the resulting initial aqueous layer was collected and lyophilised. The residue was derivatised using 2-methoxyamine hydrochloride and N-Methyl-N-(trimethylsilyl) trifluoroacetamide and subjected to gas chromatography-mass spectrometry (GC-MS) as an analytical sample. 2-Isopropylmalic acid was used as an internal standard. Additionally, an operational blank test was conducted.

The analytical equipment used included GCMS-TQ8030 (Shimadzu Corporation, Japan) and BPX5 GC columns (film thickness, 0.25 µm; length, 30 m; inner diameter, 0.25 mm; Shimadzu GC, Japan).

### Peak detection and analysis

MS-DIAL version 4.7 (http://prime.psc.riken.jp/compms/index.html) was used to analyse and prepare the peak list (peak height) under the conditions described in Supplementary Table 1. Thus, peaks that were detected in the quality control (QC) samples and whose coefficient of variation was <20% and intensity was more than twice that of the operational blank test were treated as the detected peaks.

**Table 1:**
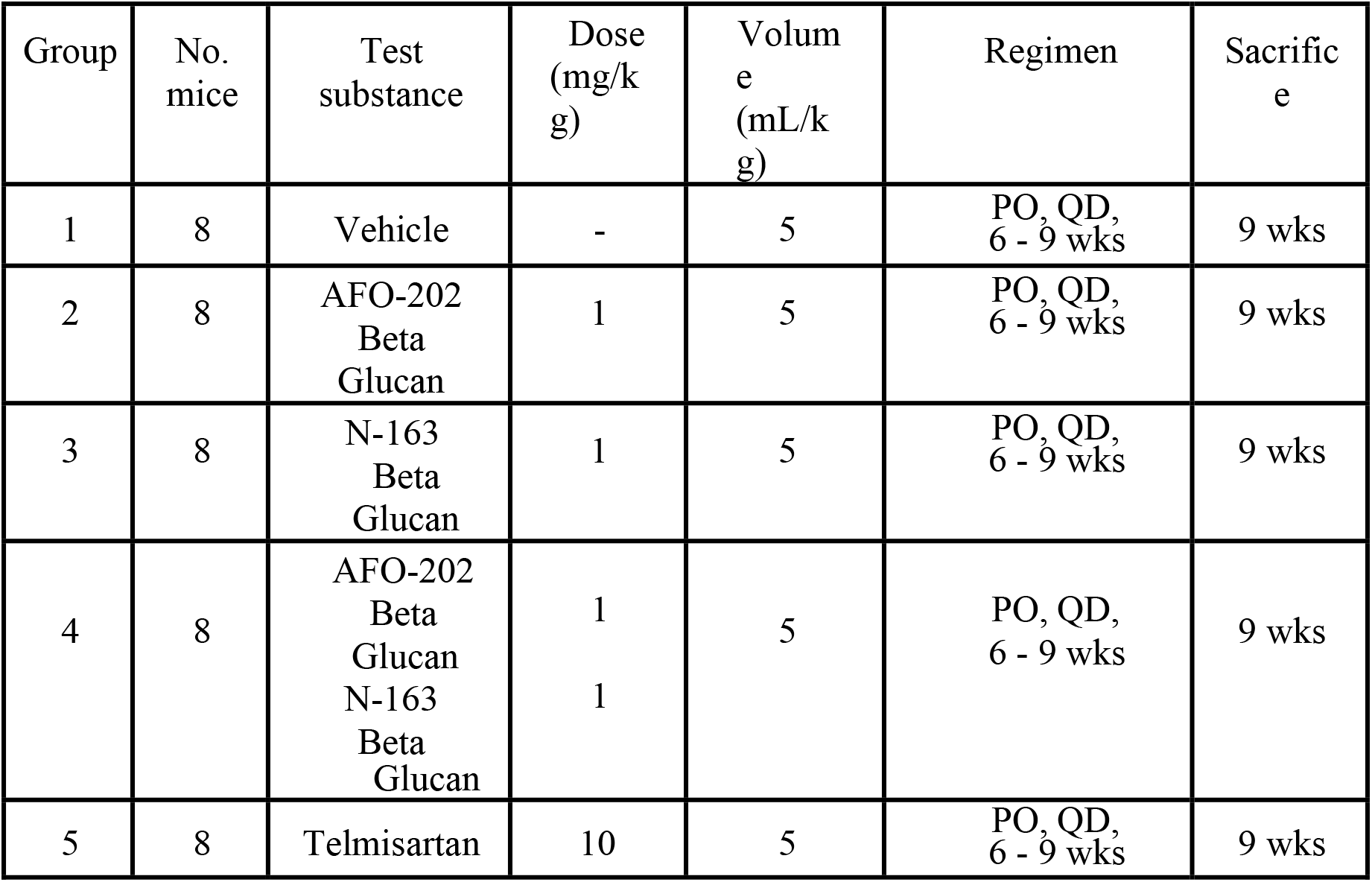
Study design and treatment schedule

### Differential abundance analysis, principal component analysis, orthogonal projections to latent structures discriminant analysis, and clustering analysis

Principal component analysis (PCA) and orthogonal partial least squares-discriminant analysis (OPLS-DA) were performed to visualise the metabolic differences among the experimental groups. For PCA, SIMCA-P+ version 17 (Umetrics) was used. The normalised peak heights of the sample-derived peaks were used for PCA using all samples and five points (F18S-12, F18S-14, F18S-16, F18S-18, and F18S-20). Transform was set to none, and scaling was set to Pareto scaling. Differential metabolites were selected according to their statistically significant variable importance in the projection (VIP) values obtained from the OPLS-DA model. R (https://www.r-project.org/) was used for hierarchical cluster analysis and generating heat maps.

### Statistical analysis

Statistical data were analysed using Microsoft Excel statistics package analysis software. Graphs were prepared using Origin Lab’s Originb 2021 software. For normally distributed variables, t-test or analysis of variance with Tukey honestly significant difference was used; P-values <0.05 were considered statistically significant. For OPLS-DA, values from two-tailed student’s t-test were applied to the normalised peak areas; metabolites with VIP values >1 and P-values <0.05 were included. The Euclidean distance and Ward method were used to analyse the heat map. The mean and variance were normalised so that the mean was 0 and the variance was 1.

## RESULTS

There were no significant differences in the mean body weight on any day during the treatment period between the control group and other treatment groups. There were no significant differences in the mean body weight on the day of sacrifice between the treatment groups.

### Effects on NASH

The effects of AFO-202 and N-163 beta-glucans on NASH were reported in our earlier paper[10] in a different set of animals subjected to the same interventions. Briefly, AFO-202 beta-glucan significantly decreased the inflammation-associated hepatic cell ballooning and steatosis, while N-163 beta-glucan significantly decreased the fibrosis and inflammation. The combination of AFO-202 and N-163 significantly decreased the NAS.[10]

### Gut microbiome analysis

The alpha-diversity indices, Simpson and Shannon indices, showed that the post-intervention gut microbial diversity was the highest in the AFO-202+N-163 group (Figure 2A and B).

**Figure 2:**
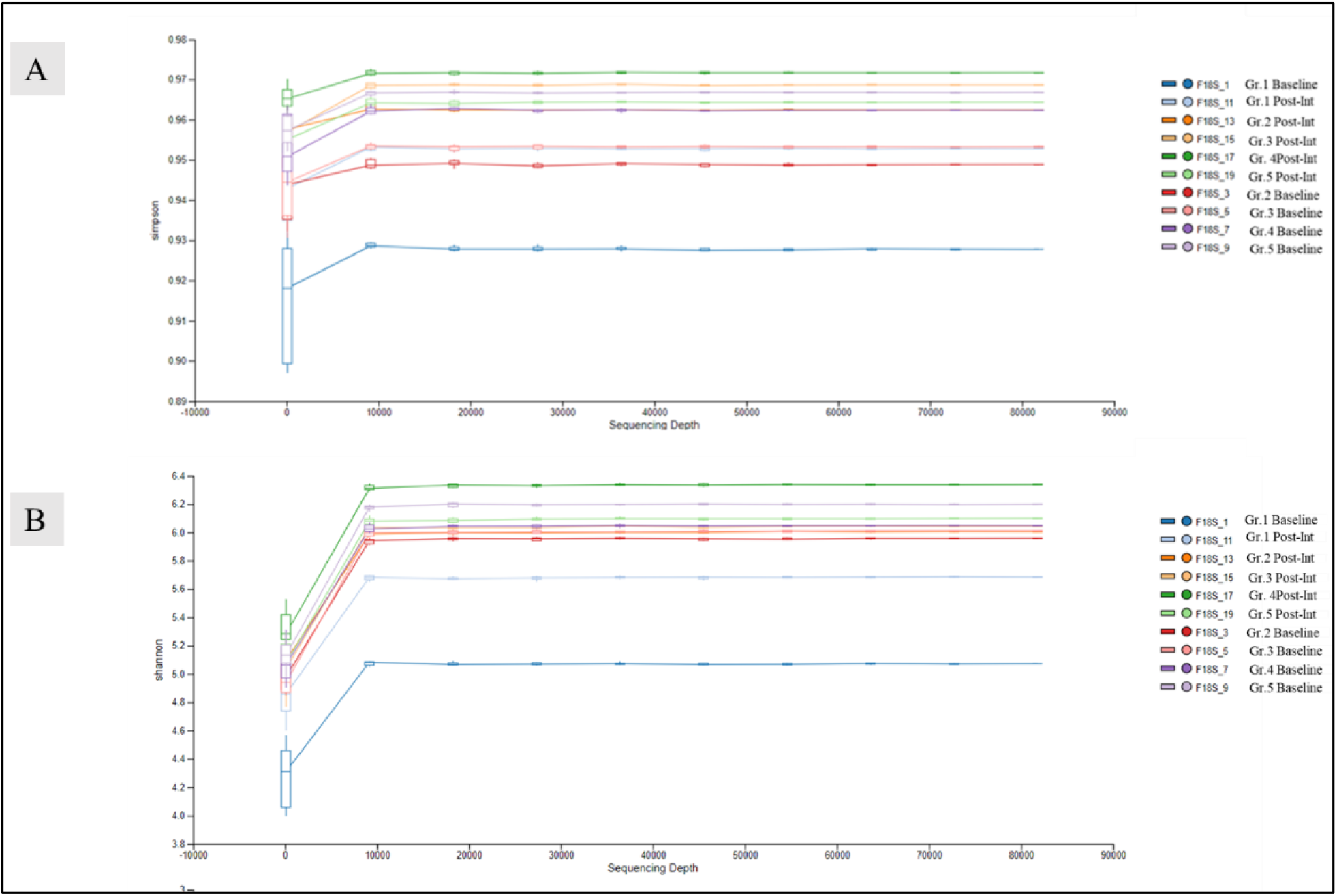
Alpha-diversity indices A. Simpson index; B. Shannon index reflecting the diversity of operational taxonomic units in samples. Both indices showed that the AFO-202+N-163 group had the highest bacterial abundance post-intervention.

With regard to taxonomic profiling, Firmicutes was the most abundant phylum, followed by Bacteroides (Figure 3).

**Figure 3:**
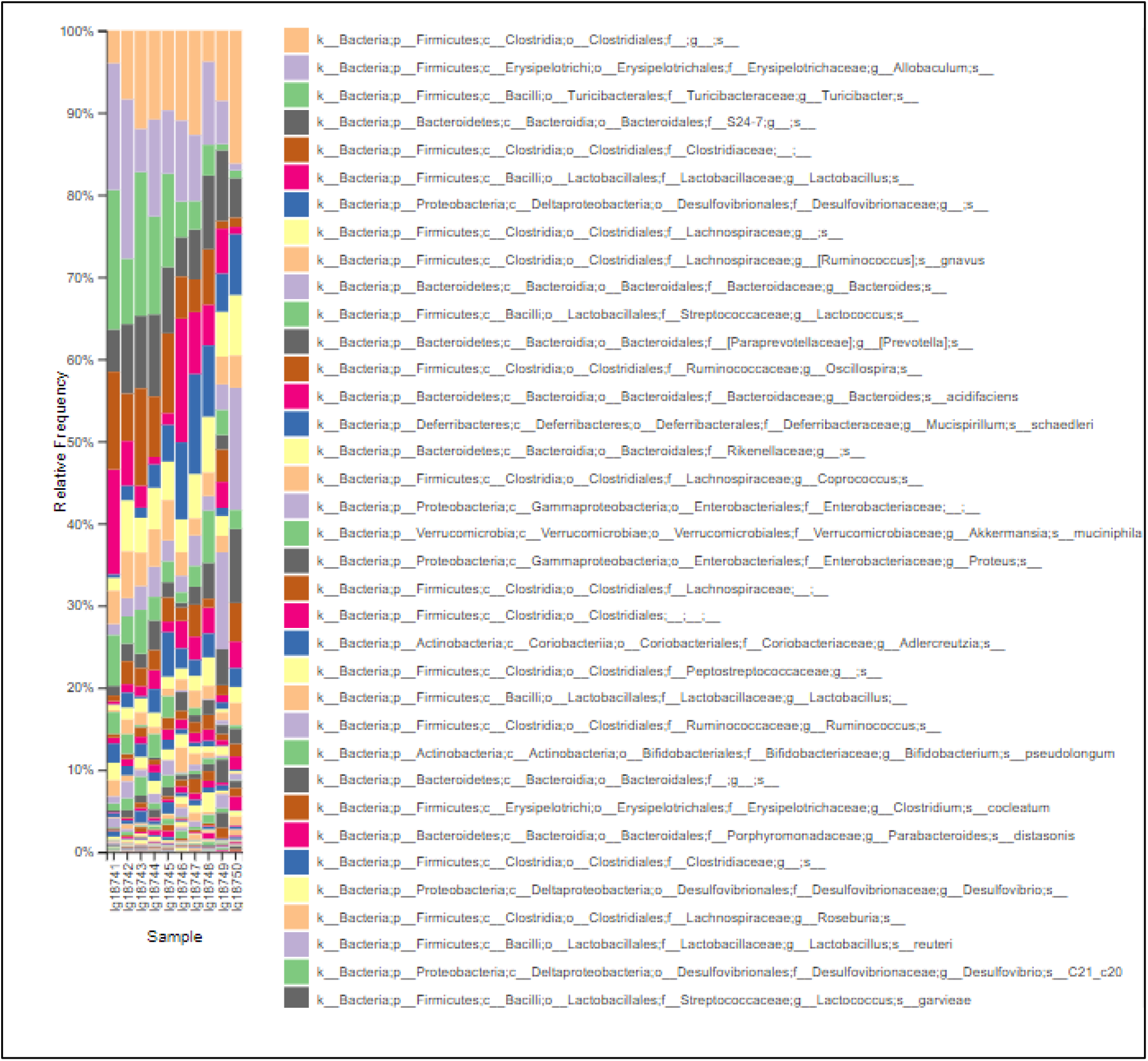
Index of the most abundant taxa across species levels

However, post-intervention, the abundance of Firmicutes decreased, while that of Bacteroides increased. This decrease and increase in the Firmicutes and Bacteroides abundance, respectively, were the highest in the AFO-202+N-163 and telmisartan groups when compared to the other groups (Supplementary Figure 1).

When individual taxa were analysed in each of the beta-glucan groups compared to the telmisartan group, the decrease in the Enterobacteria and Firmicutes abundance was the highest in the AFO-202 group (Figure 4A and B). The Turicibacter abundance was the highest in the N-163 group (Figure 4C). The Bilophila abundance increased in all groups but decreased to 0 in the N-163 group (Figure 4D). The increase in the Lactobacillus abundance was the highest in the AFO-202+N-163 group (Figure 4E). The Proteobacteria abundance decreased in the AFO-202+N-163 group but increased in the telmisartan group (Figure 4F). The decrease in Akkermansia abundance was the highest in the AFO-202+N-163 group (Figure 4G).

**Figure 4:**
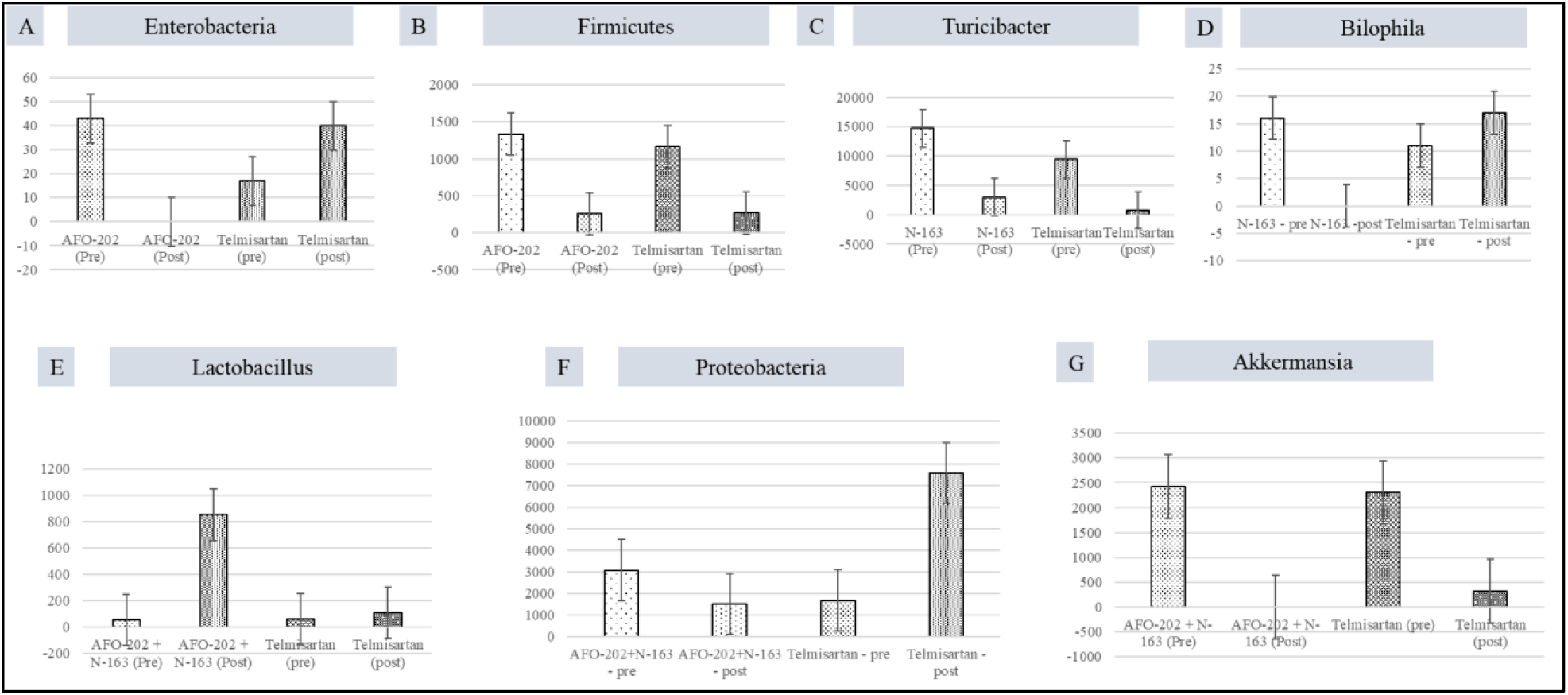
Differences between the read count of selected bacteria before and after intervention A. Enterobacteria; B. Firmicutes; C. Turicibacter; D. Bilophila; E. Lactobacillus; F. Proteobacteria; and G. Akkermansia

### Faecal metabolome analysis

The resulting score plot of the PCA using the normalised peak heights of the 10 samples (pre- and post-intervention of the five groups) is shown in Supplementary Figure 1. The contributions of the first and second principal components were 55% and 20%, respectively. PCA of the five post-intervention samples (F18S-12, F18S-14, F18S-16, F18S-18, and F18S-20) using the peak heights after normalisation and the obtained score plot are shown in Supplementary Figure 2A and B and the loading plot in Figure 2C and D. The contributions of the first and second principal components were 49% and 34%, respectively.

The peak heights of all the detected metabolite compounds after normalisation are shown in Supplementary Table 2. The number of peaks detected in the QC samples was 108, of which 53 peaks were qualitatively determined and 55 peaks remained unknown.

Differential abundance analysis and log2 fold change results are shown in Figure 5.

**Figure 5.**
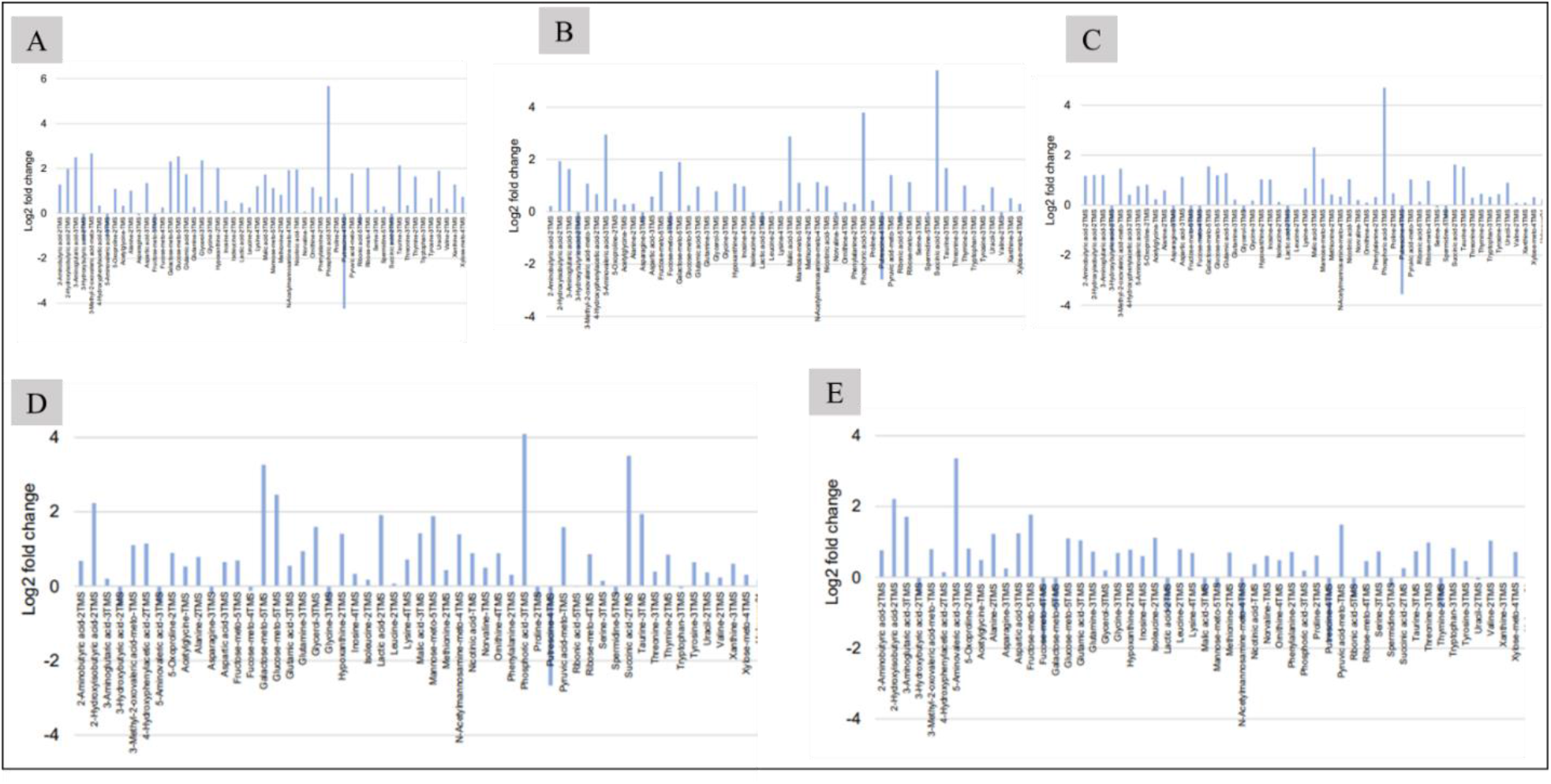
Differential abundance analysis and log2 fold change results for each group before and after intervention A. Control group; B. AFO-202 group; C. N-163 group; D. AFO-202+N-163 group; E. Telmisartan group

Score plots of PCA and compounds with VIP value ≥1 in OPLS-DA are shown in Figure 6; Supplementary Table 3 shows the compounds with VIP value ≥1 and their coefficients in OPLS-DA. PCA of the control group showed that the contribution of the first principal component axis (PC1) was 96.7% and that of the second principal component axis (PC2) was 1.5%. PCA showed that the contributions of PC1 and PC2 in the AFO-202, N-163, AFO-202+N-163, and telmisartan groups were 90.4% and 4.9%, 94.8% and 2.1%, 96.5% and 1.4%, and 95.1% and 1.9%, respectively.

**Figure 6:**
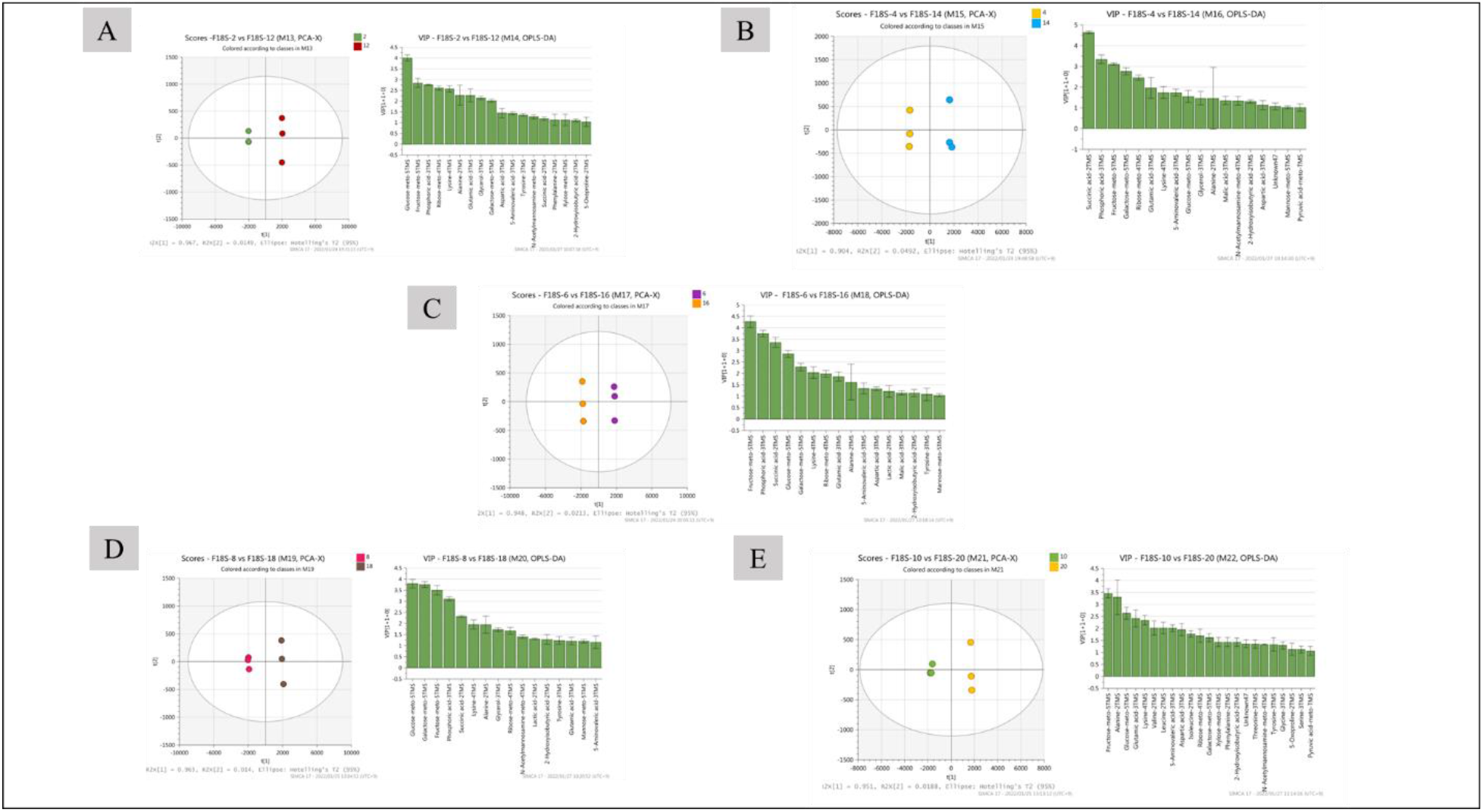
Score plots of principal component analysis and compounds with a variable importance in the projection value ≥1 in the orthogonal partial least squares-discriminant analysis of the different groups A. Control group; B. AFO-202 group; C. N-163 group; D. AFO-202+N-163 group; E. Telmisartan group

**Figure 6:**
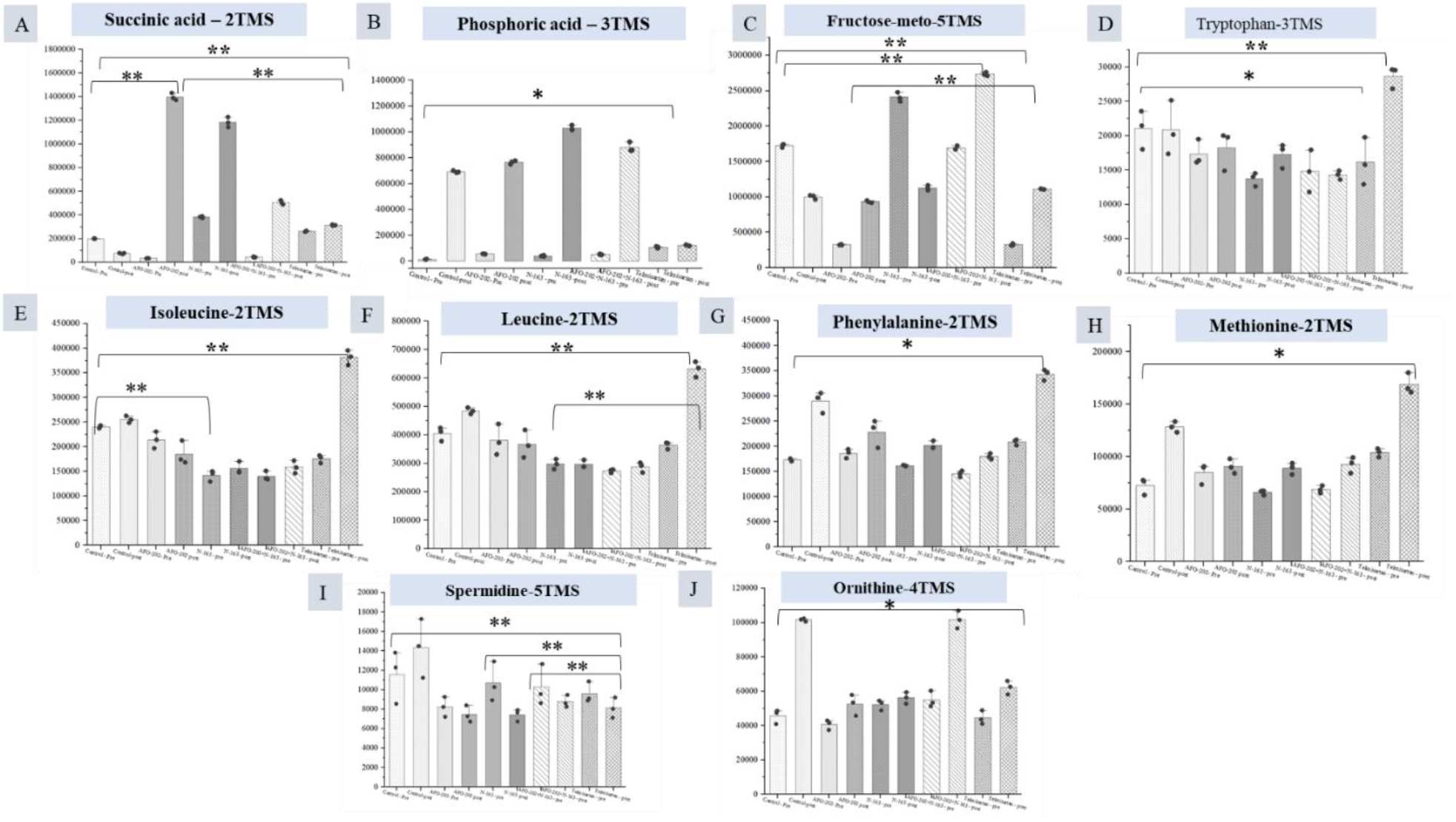
Peak heights of the detected compounds after normalisation A. Succinic acid; B. Phosphoric acid; C. Fructose; D. Tryptophan; E. Isoleucine; F. Leucine; G. Phenylalanine; H. Methionine; I. Spermidine; J. Ornithine (TMS, trimethylsilylation; meto, methoxymated derivatisation) (**, significant;*, not significant; p-value significance <0.05)

In all groups, except the telmisartan group, phosphoric acid showed the highest log2 fold increase, whereas putrescine showed the highest decrease. With respect to specific compounds, the increase in succinic acid was the highest in the AFO-202 group, with a statistical significance (P = 0.06) (Figure 6A). The increase in phosphoric acid was the highest in the N-163 group, followed by the AFO-202+N-163 and AFO-202 groups (Figure 6B), but without statistical significance (P = 0.21). The decrease in fructose was the highest in the AFO-202 group (P = 0.0007) (Figure 6C). Tryptophan decreased in the AFO-202+N-163 group, but not significantly (P = 0.99); however, it increased in the other groups (Figure 6D). The decrease in isoleucine and leucine was the highest in the N-163 group, with statistical significance (P = 0.004 and 0.012, respectively) (Figure 6E and F). The decrease in phenylalanine was the highest in the AFO-202+N-163 group, without statistical significance (P = 0.18) (Figure 6G). Methionine increased in all the groups but without statistical significance (P = 0.14) (Figure 6H). The decrease in spermidine was the highest in the N-163 group, with statistical significance (P = 0.012) (Figure 6I). The increase in ornithine was the highest in the AFO-202+N-163 group (Figure 6J). Euclidean distance hierarchical clustering analysis demonstrated that the different intensity levels of characteristic metabolites also matched the aforementioned findings (Figure 7 and Supplementary Figure 3).

**Figure 7:**
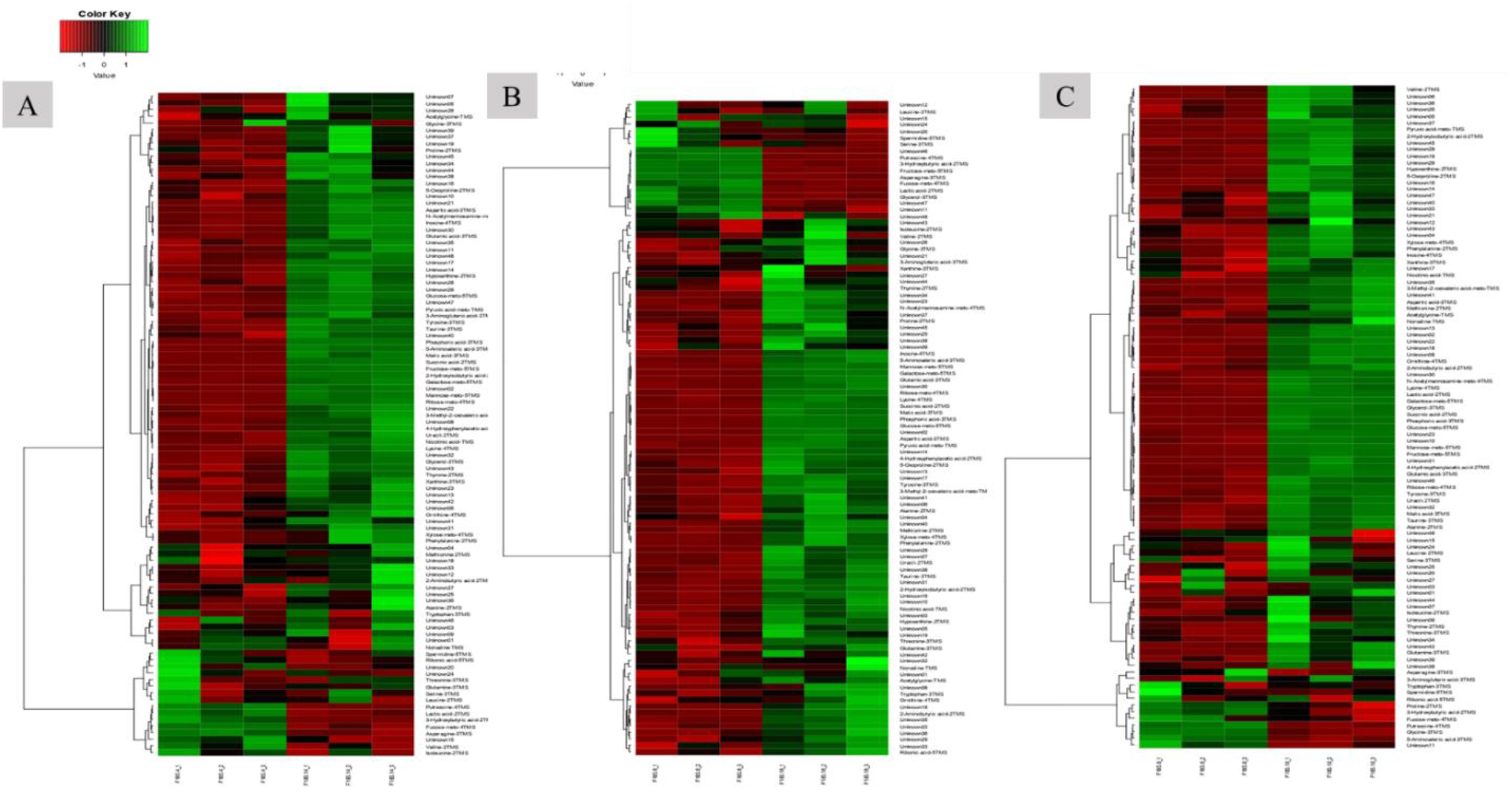
Euclidean distance hierarchical clustering analysis demonstrating the different intensity levels of characteristic metabolites in the treatment groups A. AFO-202 group; B. N-163 group; C. AFO-202+N-163 (Comparison images of the telmisartan and control groups are available in Supplementary Figure 3.)

## DISCUSSION

This is the first study to investigate the influence of beta-glucans on the profiles of the faecal gut microbiome and metabolome in a NASH murine model. This study assessed two different beta-glucans produced by different strains of the same species of black yeast, A. pullulans. Beta-glucans are obtained from different sources, and their functionality depends on the source and extraction/purification processes.[14] The beta-glucans described in this study, AFO-202 and N-163 strains of A. pullulans black yeast, are unique as they are produced as an exopolysaccharide without the need for extraction/purification; hence, their biological actions are superior to those of the other strains.[15] Furthermore, both the beta-glucans have the same chemical formula but different structural formulas, and hence exert diverse biological actions. AFO-202 beta-glucan has beneficial metabolic benefits as it regularises blood glucose levels[5] and enhances immunity in immune-related infections such as COVID-19[11, 12]. Moreover, it produces positive effects on melatonin and alpha-synuclein neurotransmitters and sleep and behaviour in neurodevelopmental disorders such as ASD.[7, 8] In a previous NASH animal study, AFO-202 beta-glucan significantly decreased the inflammation-associated hepatic cell ballooning and steatosis.[10] The N-163 beta-glucan has immunomodulatory benefits in terms of regulating dyslipidaemia, which is evident from the balance in the levels of non-esterified fatty acids[16] and decrease in fibrosis and inflammation in NASH.[10] The combination of AFO-202 and N-163 decreased pro-inflammatory markers and increased anti-inflammatory markers in healthy human volunteers,[17] decreased the NAS in the NASH model,[10] and significantly controlled immune-mediated dysregulated levels of interleukin-6, C-reactive protein, and ferritin in COVID-19 patients.[11, 12] In a study on gut microbiome analysis in ASD patients, AFO-202 showed efficient control of Enterobacteria as well as beneficial reconstitution of the gut microbiome with positive effects on ASD.[13] This study aimed to evaluate the benefits of AFO-202 and N-163 individually and in combination in an animal model of NASH.

We used the STAM of NASH.[10, 18, 19] In this model, HFD-fed mice were allowed to develop liver steatosis by administering streptozotocin solution 2 days after birth. This model recapitulates most of the features of the metabolic syndrome of NASH that occur in humans, and the HFD leads to diabetes, dyslipidaemia, and liver steatosis. Therefore, the gut microbiome profiles and faecal metabolite profiles present at baseline recapitulate those of metabolic syndrome.[20, 21] In time, this will produce pathophysiological problems in different organ systems of the body, including the heart, liver, and kidneys, as well as immune-metabolic interactions, leading to a decline in the immune system with aging and its associated complications. Therefore, this study could serve as a forerunner to study the effects of beta-glucans on the different aspects of metabolic syndrome-associated pathologies as well as conditions associated with immune-metabolic interactions, including neurological disorders wherein immune-metabolic interactions have profound implications.[20]

### Relevance of metabolome and microbiome

An abundance of bacterial species such as Proteobacteria, Enterobacteria, and Escherichia coli has been reported in humans with NAFLD. Greater abundance of Prevotella species has been reported in obese children with NAFLD.[21, 22] In this study, there was a decrease in the Enterobacteria abundance with AFO-202 and a significant decrease in the Prevotella abundance with AFO-202+N-163. In terms of faecal metabolites, an increase in tryptophan was observed in the N-163 group, but it was not more than that in the telmisartan group. In NAFLD, tryptophan metabolism is disturbed, and supplementation of tryptophan was reportedly beneficial because it increased the intestinal integrity and improved liver steatosis and function in an NAFLD mouse model.[22] Decreased production of butyrate increases intestinal inflammation, gut permeability, endotoxaemia, and systemic inflammation. An increased abundance of 2-hydroxyisobutyric acid was observed in the AFO-202+N-163 group (Figure 8 and Supplementary Table 2).

### Potential in neurological illnesses

#### Neurodevelopmental and neurodegenerative disorders

In addition to the influence of beta-glucans on the profiles of the faecal gut microbiome and metabolome in a NASH murine model, we previously reported a decrease in the abundance of Enterobacteria, E. coli, Akkermansia muciniphila CAG:154, Blautia spp., Coprobacillus spp., and Clostridium bolteae CAG:59. Furthermore, there was an increase in the abundance of Faecalibacterium prausnitzii and Prevotella copri after AFO-202 administration in children with ASD.[13] In this study, the decrease in Enterobacteria abundance was the highest in the AFO-202 group (Figure 5A). Succinic acid, which is reportedly low in ASD individuals,[23] was found to be the highest in the AFO-202 group (Figure 7A). In several neurodegenerative diseases, such as PD, the levels of amino acids such as isoleucine, leucine, and phenylalanine have been found to be high in the faecal metabolome.[24] In this study, the decrease in the levels of these amino acids was the highest in the AFO-202 group (Figure 7 E–G).

#### Neuroinflammatory disorders

Turicibacter has been associated with inflammatory conditions, such as inflammatory bowel disease, and other chronic immune-mediated inflammatory diseases, such as MS, due to its correlation with the expression of tumour necrosis factor.[25] In this study, the N-163 and telmisartan groups showed a decrease in the Turicibacter abundance (Figure 5B). Similarly, a significant increase in the relative abundance of Desulfovibrionaceae (Bilophila) was reported in early-onset paediatric MS.[26] In this study, the N-163 group showed a reduction in the abundance of Bilophila, whereas the telmisartan group showed an increase after intervention (Figure 5E). Corroborating this finding, inflammatory conditions present high amounts of sulphur-containing metabolites, such as methionine,[27] whose increase in abundance was the least in the N-163 group (Figure 7H). The presence of A. muciniphila, a mucosal-dwelling anaerobe, is a double-edged sword.[28] It was previously reported that its abundance reduced in various metabolic disorders, including obesity, dyslipidaemia, and type 2 diabetes, which fuelled the development of Akkermansia-based probiotic therapies to combat metabolic disorders.[29] However, recent evidence suggests that an increase in Akkermansia abundance has been reported in PD and MS patients.[30] In this study, there was a significant decrease in the Akkermansia abundance in the AFO-202+N-163 group (Figure 5G).

### Other implications

In overweight/obese humans, low faecal bacterial diversity is reportedly associated with a marked increase in fat tissue dyslipidaemia, impaired glucose homeostasis, and increased incidence of low-grade inflammation.[31] In this study, the bacterial diversity increased after the intervention, especially in the AFO-202+N-163 group, which showed the highest diversity in the Shannon and Simpson indices (Figure 2). Most studies report an increase in Firmicutes and decrease in Bacteroides abundance to be directly proportional to body weight gain.[31, 32] In this study, there was a clear decrease in Firmicutes and increase in Bacteroides abundance in all the groups post-intervention, but the highest change was in the AFO-202+N-163 and telmisartan groups (Figures 3 and 4). Spermidine is a metabolite that has been associated with inflammation and cancer.[33] The decrease in spermidine was the highest in the N-163 group (Figure 7I). Ornithine levels are usually low in colorectal cancer patients.[34] The increase in ornithine levels was the highest in the AFO-202+N-163 group in this study. Lactobacillus is a common probiotic used as prophylaxis and treatment in chronic conditions such as cancer[35] as well as for promoting better health. The increase in Lactobacillus abundance was the highest in the AFO-202+N-163 group (Figure 5D). Steroids are common immunosuppressants used to treat chronic autoimmune conditions as well as organ transplant recipients. The use of steroids has been reported to cause an increase in E. Coli and Enterococcus and a decrease in Bacteroides abundance.[36] In this study, the control of Enterobacteria and an increase in Bacteroides abundance with AFO-202, N-163, and their combination makes them worthy adjuncts to medications such as steroids.

## CONCLUSION

Two strains of black yeast A. pullulans, AFO-202 and N-163, produce beta glucans that increase the gut microbial diversity, control harmful bacteria, promote healthy bacteria, and induce beneficial changes in the faecal metabolites, all indicative of a healthy profile, both individually and in combination in the NASH animal model. The results of this study support the use of AFO-202 as a potentially effective and adjunct treatment in several neurodevelopmental conditions such as ASD and neurodegenerative conditions such as PD. N-163 can help to control inflammation and chronic immune-associated inflammatory conditions such as MS. The combination of AFO-202 and N-163 could help to preserve the overall health, serving as a preventive agent against chronic inflammatory and immune-dysregulated conditions such as cancer. However, further trials are warranted to determine if the use of beta-glucans would be useful in the treatment of other chronic human inflammatory conditions.

## Acknowledgments

The authors would like to dedicate this paper to the memory of Mr. Takashi Onaka, who passed away on the 1st of June, 2022 at the age of 90 years, who played an instrumental role in successfully culturing and industrial scale up of AFO-202 and N-163 strains of Aureobasidium pullulans after their isolation by Prof. Noboru Fujii, producing the novel beta glucans described in this study. They thank, Mr. Yasushi Onaka, Mr. Masato Onaka, Mr. Yasunori Ikeue, Dr. Mitsuru Nagataki and Mr. Ken Sakanishi of Sophy Inc., Mr. Yoshio Morozumi and Ms. Yoshiko Amikura of GN Corporation, Japan and Loyola-ICAM College of Engineering and Technology (LICET) for their support to our research work.

## Availability of data and material

All data generated or analysed during this study are included in the article itself.

## Ethics Approval

Protocol approvals were obtained from SMC Laboratories, Japan’s IACUC (Study reference no: SLMN070-2108-9)

All animals used in this study were cared for under the following guidelines: Act on Welfare and Management of Animals (Ministry of the Environment, Japan, Act No. 105 of October 1, 1973), standards relating to the care and management of laboratory animals and relief of pain (Notice No.88 of the Ministry of the Environment, Japan, April 28, 2006) and the guidelines for proper conduct of animal experiments (Science Council of Japan, June 1, 2006).

## References

1. Valdes AM, Walter J, Segal E, Spector TD. Role of the gut microbiota in nutrition and health. BMJ. 2018 Jun 13;361:k2179.

2. Ghaisas S, Maher J, Kanthasamy A. Gut microbiome in health and disease: Linking the microbiome-gut-brain axis and environmental factors in the pathogenesis of systemic and neurodegenerative diseases. Pharmacol Ther. 2016 Feb;158:52–62.

3. Shreiner AB, Kao JY, Young VB. The gut microbiome in health and in disease. Curr Opin Gastroenterol. 2015 Jan;31(1):69–75. doi: 10.1097/MOG.0000000000000139. PMID: 25394236; PMCID: PMC4290017.

4. Zierer J, Jackson MA, Kastenmüller G, Mangino M, Long T, Telenti A, Mohney RP, Small KS, Bell JT, Steves CJ, Valdes AM, Spector TD, Menni C. The fecal metabolome as a functional readout of the gut microbiome. Nat Genet. 2018 Jun;50(6):790–795.

5. Dedeepiya V, Sivaraman G,Venkatesh A, Preethy S, Abraham S. Potential Effects of Nichi Glucan as a Food Supplement for Diabetes Mellitus and Hyperlipidemia; Preliminary Findings from the Study on Three Patients from India. Case Reports in Medicine 2012 (2012), Article ID 895370

6. Ganesh JS, Rao YY, Ravikumar R, Jayakrishnan AG, Iwasaki M, Preethy S, Abraham S. Beneficial effects of Black yeast derived 1-3, 1-6 beta glucan-Nichi Glucan in a dyslipidemic individual of Indian origin - A case report. J Diet Suppl. 2014;11(1):1–6.

7. Raghavan K, Dedeepiya VD, Ikewaki N, Sonoda T, Iwasaki M, Preethy S, Abraham SJK. Improvement of behavioural pattern and alpha-synuclein levels in autism spectrum disorder after consumption of a beta-glucan food supplement in a randomized, parallel-group pilot clinical study. BMJ Neurology Open (In print)

8. Raghavan K, Dedeepiya VD, Kandaswamy R, Balamurugan M, Ikewaki N, Sonoda T, Kurosawa G, Iwasaki M, Preethy S, Abraham SJK. Improvement of sleep patterns and serum melatonin levels in children with autism spectrum disorders after consumption of beta-1,3/1,6-glucan in a pilot clinical study. Research Square rs.3.rs-701988/v1; doi: 10.21203/rs.3.rs-701988/v1

9. Raghavan K, Dedeepiya VD, Srinivasan S, Pushkala S, Subramanian S, Ikewaki N, Iwasaki M, Senthilkumar R, Preethy S, Abraham S. Disease-modifying immune-modulatory effects of the N-163 strain of Aureobasidium pullulans-produced 1,3-1,6 Beta glucans in young boys with Duchenne muscular dystrophy: Results of an open-label, prospective, randomized, comparative clinical study. medRxiv 2021.12.13.21267706

10. Ikewaki N, Kurosawa G, Iwasaki M, Preethy S, Dedeepiya VD, Vaddi S, Senthilkumar R, Levy GA, Abraham SJK. Hepatoprotective effects of Aureobasidium pullulans derived Beta 1,3-1,6 biological response modifier glucans in a STAM-animal model of non-alcoholic steatohepatitis. bioRxiv 2021.07.08.451700; doi: 10.1101/2021.07.08.451700

11. Pushkala S, Seshayyan S, Theranirajan E, Sudhakar D, Raghavan K, Dedeepiya VD, Ikewaki N,Iwasaki M, Preethy S, Abraham S. Efficient control of IL-6, CRP and Ferritin in Covid-19 patients with two variants of Beta-1,3-1,6 glucans in combination, within 15 days in an open-label prospective clinical trial. medRxiv 2021.12.14.21267778; doi: 10.1101/2021.12.14.21267778

12. Raghavan K, Dedeepiya VD, Suryaprakash V, Rao KS, Ikewaki N, Sonoda T, Levy GA, Iwasaki M, Senthilkumar R, Preethy S, Abraham SJK. Beneficial Effects of novel aureobasidium pullulans strains produced beta-1,3-1,6 glucans on interleukin-6 and D-Dimer levels in COVID-19 patients; results of a randomized multiple-arm pilot clinical study. Biomedicine and Pharmacotherapy 2021. sciencedirect. https://doi.org/10.1016/j.biopha.2021.112243

13. Raghavan K, Dedeepiya VD, Yamamoto N, Ikewaki N, Sonoda T, Kurosawa G, Iwasaki M, Kandaswamy R, Senthilkumar R, Preethy S, Abraham SJK. Beneficial reconstitution of gut microbiota and control of alpha-synuclein and curli-amyloids-producing enterobacteria, by beta 1,3-1,6 glucans in a clinical pilot study of autism and potentials in neurodegenerative diseases. medRxiv 2021.10.26.21265505; doi: 10.1101/2021.10.26.21265505

14. Bashir KMI, Choi JS. Clinical and Physiological Perspectives of β-Glucans: The Past, Present, and Future. Int J Mol Sci. 2017 Sep 5;18(9):1906.

15. Ikewaki N, Fujii N, Onaka T, Ikewaki S, Inoko H. Immunological actions of Sophy betaglucan (beta-1,3-1,6 glucan), currently available commercially as a health food supplement. Microbiol Immunol. 2007;51(9):861–73.

16. Ikewaki N, Onaka T, Ikeue Y, Nagataki M, Kurosawa G, Dedeepiya VD, Rajmohan M, Vaddi S, Senthilkumar R, Preethy S, Abraham SJK. Beneficial effects of the AFO-202 and N-163 strains of Aureobasidium pullulans produced 1,3-1,6 beta glucans on non-esterified fatty acid levels in obese diabetic KKAy mice: A comparative study. bioRxiv 2021.07.22.453362; doi: 10.1101/2021.07.22.453362

17. Ikewaki N, Sonoda T, Kurosawa G, Iwasaki M, Dedeepiya VD, Senthilkumar R, Preethy S, Abraham SJK. Immune and metabolic beneficial effects of Beta 1,3-1,6 glucans produced by two novel strains of Aureobasidium pullulans in healthy middle-aged Japanese men: An exploratory study. medRxiv 2021.08.05.21261640; doi: 10.1101/2021.08.05.21261640

18. STAM Model. https://www.smccro-lab.com/service/service_disease_area/stam.html

19. Nakashima A, Sugimoto R, Suzuki K, Shirakata Y, Hashiguchi T, Yoshida C, Nakano Y. Anti-fibrotic activity of Euglena gracilis and paramylon in a mouse model of non-alcoholic steatohepatitis. Food Sci Nutr. 2018;7(1):139–147

20. Dantzer R. Neuroimmune Interactions: From the Brain to the Immune System and Vice Versa. Physiol Rev. 2018 Jan 1;98(1):477–504.

21. Kolodziejczyk AA, Zheng D, Shibolet O, Elinav E. The role of the microbiome in NAFLD and NASH. EMBO Mol Med. 2019 Feb;11(2):e9302. doi: 10.15252/emmm.201809302. PMID: 30591521; PMCID: PMC6365925.

22. Chen J, Vitetta L. Gut Microbiota Metabolites in NAFLD Pathogenesis and Therapeutic Implications. Int J Mol Sci. 2020 Jul 23;21(15):5214. doi: 10.3390/ijms21155214. PMID: 32717871; PMCID: PMC7432372.

23. Kang DW, Adams JB, Vargason T, Santiago M, Hahn J, Krajmalnik-Brown R. Distinct Fecal and Plasma Metabolites in Children with Autism Spectrum Disorders and Their Modulation after Microbiota Transfer Therapy. mSphere. 2020 Oct 21;5(5):e00314–20. doi: 10.1128/mSphere.00314-20. PMID: 33087514; PMCID: PMC7580952.

24. Vascellari S, Palmas V, Melis M, Pisanu S, Cusano R, Uva P, Perra D, Madau V, Sarchioto M, Oppo V, Simola N, Morelli M, Santoru ML, Atzori L, Melis M, Cossu G, Manzin A. Gut Microbiota and Metabolome Alterations Associated with Parkinson’s Disease. mSystems. 2020 Sep 15;5(5):e00561–20. doi: 10.1128/mSystems.00561-20. PMID: 32934117; PMCID: PMC7498685.

25. Bernstein CN, Forbes JD. Gut Microbiome in Inflammatory Bowel Disease and Other Chronic Immune-Mediated Inflammatory Diseases. Inflamm Intest Dis. 2017 Nov;2(2):116–123. doi: 10.1159/000481401. Epub 2017 Oct 20. PMID: 30018962; PMCID: PMC5988152.

26. Tremlett H, Fadrosh DW, Faruqi AA, Zhu F, Hart J, Roalstad S, Graves J, Lynch S, Waubant E; US Network of Pediatric MS Centers. Gut microbiota in early pediatric multiple sclerosis: a case-control study. Eur J Neurol. 2016 Aug;23(8):1308–1321.

27. Walker A, Schmitt-Kopplin P. The role of fecal sulfur metabolome in inflammatory bowel diseases. Int J Med Microbiol. 2021 Jul;311(5):151513.

28. Cirstea M, Radisavljevic N, Finlay BB. Good Bug, Bad Bug: Breaking through Microbial Stereotypes. Cell Host Microbe. 2018 Jan 10;23(1):10–13.

29. Berer K, Gerdes LA, Cekanaviciute E, Jia X, Xiao L, Xia Z, Liu C, Klotz L, Stauffer U, Baranzini SE, Kümpfel T, Hohlfeld R, Krishnamoorthy G, Wekerle H. Gut microbiota from multiple sclerosis patients enables spontaneous autoimmune encephalomyelitis in mice. Proc Natl Acad Sci U S A. 2017 Oct 3;114(40):10719–10724. doi: 10.1073/pnas.1711233114. Epub 2017 Sep 11. PMID: 28893994; PMCID: PMC5635914.

30. Heintz-Buschart A, Pandey U, Wicke T, Sixel-Döring F, Janzen A, Sittig-Wiegand E, Trenkwalder C, Oertel WH, Mollenhauer B, Wilmes P. The nasal and gut microbiome in Parkinson’s disease and idiopathic rapid eye movement sleep behavior disorder. Mov Disord. 2018 Jan;33(1):88–98. doi: 10.1002/mds.27105. Epub 2017 Aug 26. PMID: 28843021; PMCID: PMC5811909.

31. Davis CD. The Gut Microbiome and Its Role in Obesity. Nutr Today. 2016 Jul-Aug;51(4):167–174. doi: 10.1097/NT.0000000000000167. PMID: 27795585; PMCID: PMC5082693.

32. Jumpertz R, Le DS, Turnbaugh PJ, et al. Energy-balance studies reveal associations between gut microbes, caloric load, and nutrient absorption in humans. Am. J. Clin. Nutr. 2011;94:58–65.

33. Yoshimoto S, Mitsuyama E, Yoshida K, Odamaki T, Xiao JZ. Enriched metabolites that potentially promote age-associated diseases in subjects with an elderly-type gut microbiota. Gut Microbes. 2021 Jan-Dec;13(1):1–11.

34. Le Gall G, Guttula K, Kellingray L, Tett AJ, Ten Hoopen R, Kemsley EK, Savva GM, Ibrahim A, Narbad A. Metabolite quantification of faecal extracts from colorectal cancer patients and healthy controls. Oncotarget. 2018 Sep 7;9(70):33278–33289. doi: 10.18632/oncotarget.26022. Erratum in: Oncotarget. 2019 Feb 26;10(17):1660. PMID: 30279959; PMCID: PMC6161785.

35. Lu K, Dong S, Wu X, Jin R, Chen H. Probiotics in Cancer. Front Oncol. 2021 Mar 12;11:638148.

36. Gibson CM, Childs-Kean LM, Naziruddin Z, Howell CK. The alteration of the gut microbiome by immunosuppressive agents used in solid organ transplantation. Transpl Infect Dis. 2021 Feb;23(1):e13397.

