## Supplemental Figure for "Two unique biological response-modifier glucans beneficially regulating gut microbiota and faecal metabolome in a non-alcoholic steatohepatitis animal model, with potential for applications in human health and disease"

**Supplementary material**

**Supplemental Figures**

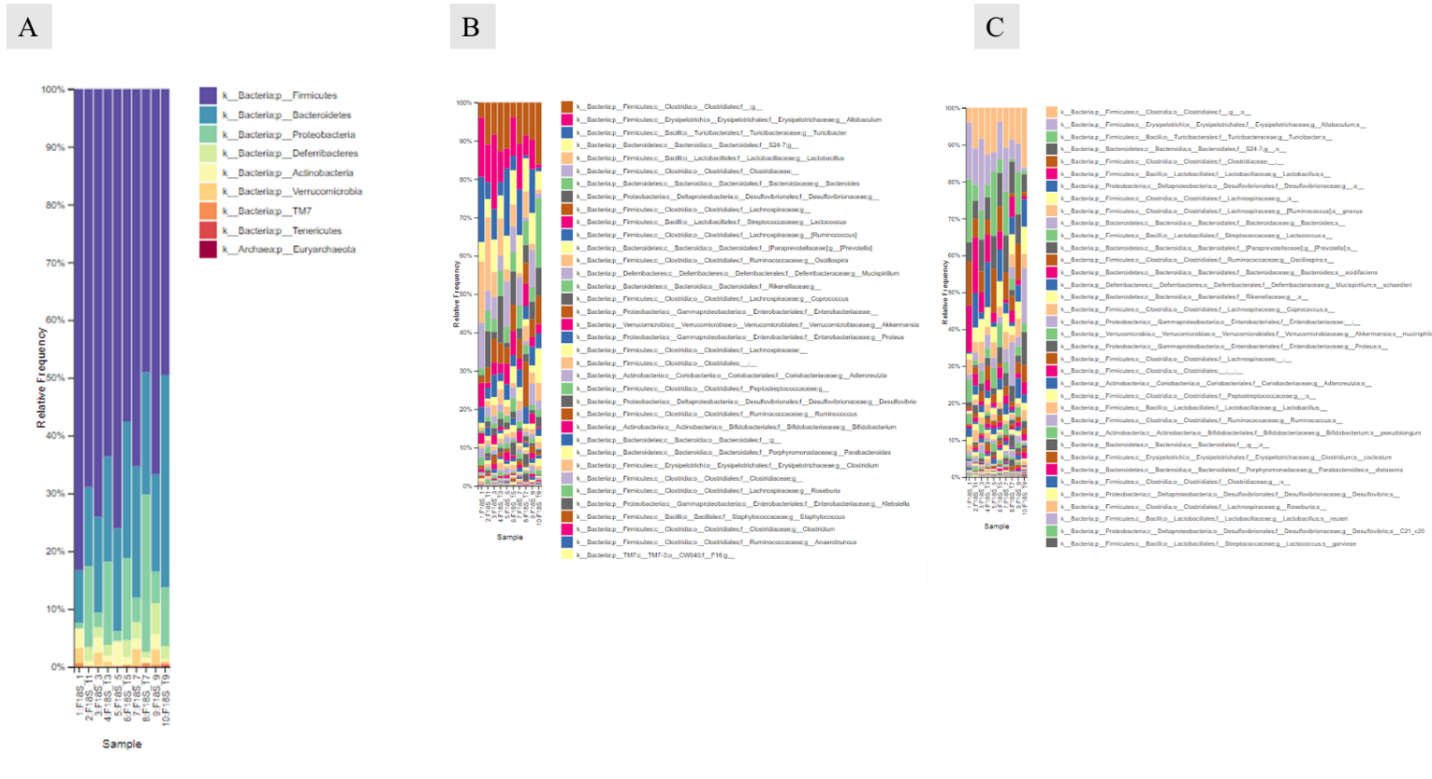

Supplementary Figure 1: Most abundant taxa at the phylum (A), genus (B), and species (C) levels in the different study groups

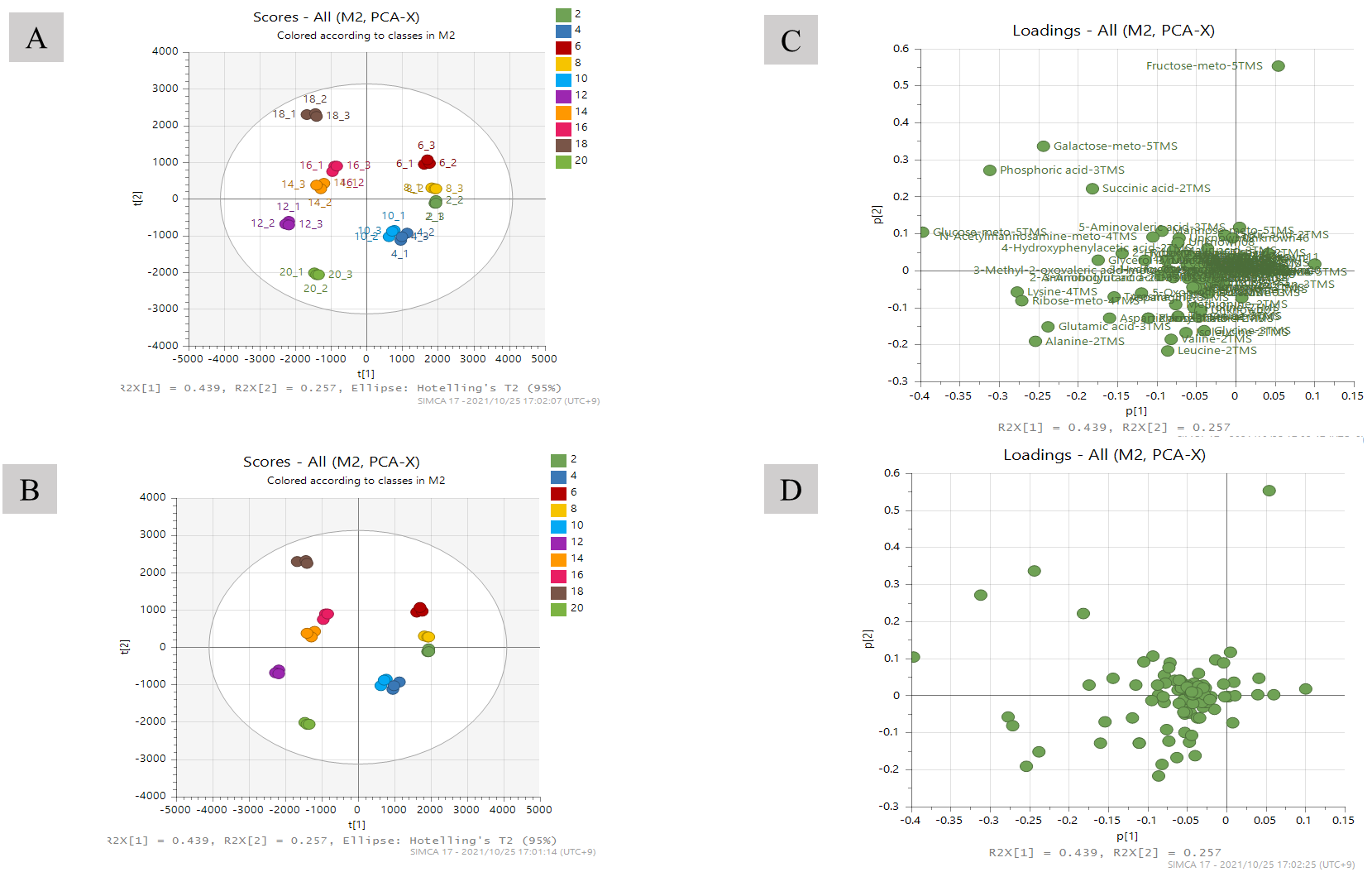
Supplementary Figure 2: Principal component analysis of all ten samples (five groups: pre- and post-intervention)

A, B. Score plot; C, D. Loading plot

Control/vehicle group: Baseline, 2; Post-intervention, 12; AFO-202 group: Baseline, 4; Post-intervention, 14; N-163 group: Baseline, 6; Post-intervention, 16; AFO-202+N-163 group: Baseline, 8; Post-intervention, F18S-18; Telmisartan group: Baseline, 8; Post-intervention, 18

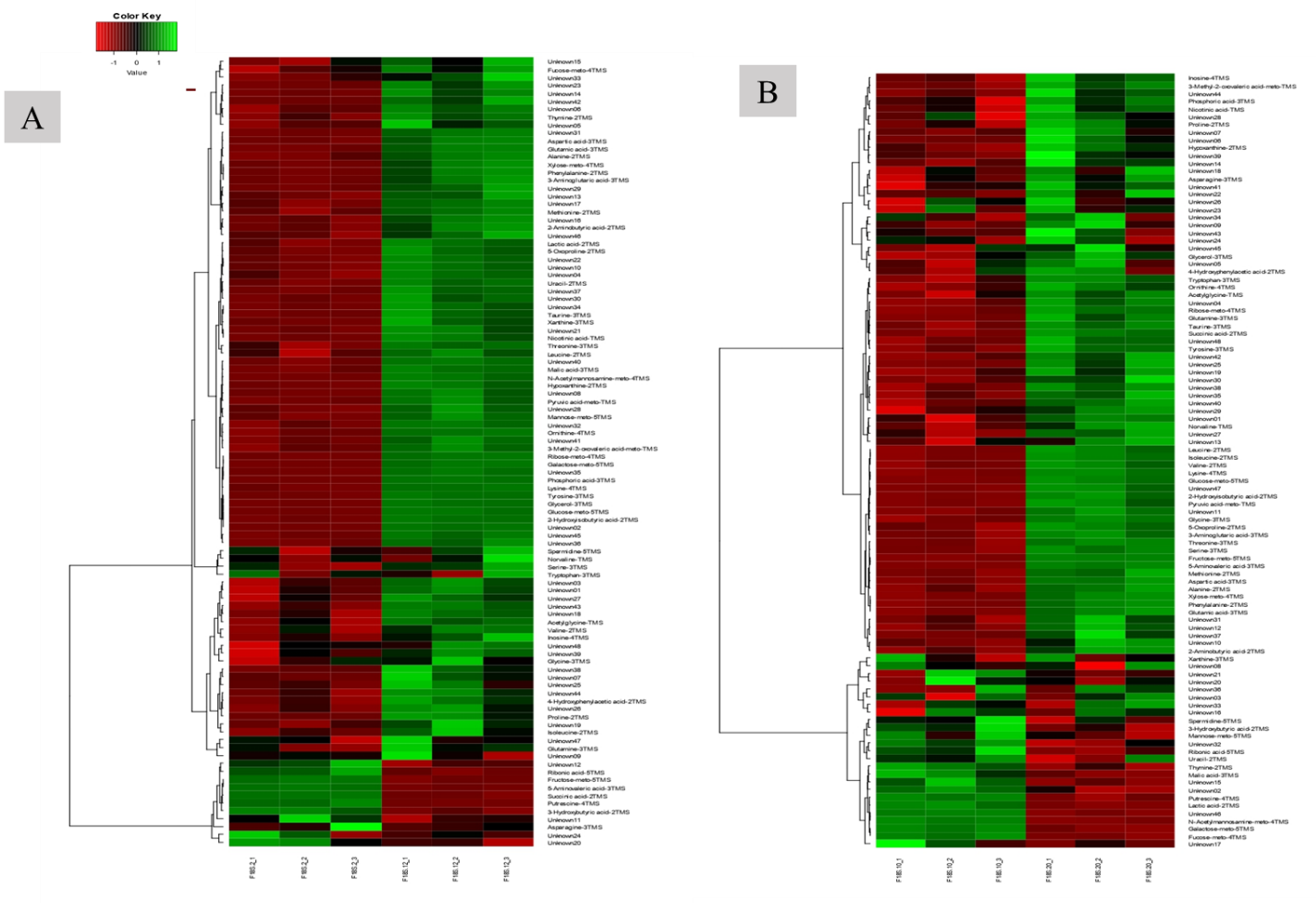
Supplementary Figure 3: Euclidean distance hierarchical clustering analysis demonstrating the different intensity levels of characteristic metabolites in the control group (A) and telmisartan group (B)
