## Supplementary Table for "Two unique biological response-modifier glucans beneficially regulating gut microbiota and faecal metabolome in a non-alcoholic steatohepatitis animal model, with potential for applications in human health and disease"

**Supplementary Tables**

**Supplementary Table 1: Abundance of all the bacteria analysed at baseline and post-intervention**

| **Order** | **Family** | **Genus** | **Species** | **Control (Pre)** | **Control (Post)** | **AFO-202 (Pre)** | **AFO-202 (Post)** | **N-163 (Pre)** | **N-163 (Post)** | **AFO-202+N-163 (Pre)** | **AFO-202+N-163 (Post)** | **Telmisartan (Pre)** | **Telmisartan (Post)** |
| --- | --- | --- | --- | --- | --- | --- | --- | --- | --- | --- | --- | --- | --- |
| 'Erysipelotrichales | 'Erysipelotrichaceae | 'Allobaculum | '_ | 12083 | 8007 | 15713 | 6542 | 4491 | 7214 | 9897 | 4360 | 6375 | 622 |
| 'Turicibacterales | 'Turicibacteraceae | 'Turicibacter | '_ | 12538 | 3346 | 6159 | 2749 | 14702 | 2976 | 9032 | 630 | 9408 | 786 |
| 'Desulfovibrionales | 'Desulfovibrionaceae | '_ | '_ | 306 | 8016 | 1398 | 10096 | 1070 | 7145 | 2351 | 3790 | 3882 | 6012 |
| 'Clostridiales | 'Clostridiaceae | 'Other | 'Other | 7455 | 3479 | 3689 | 2806 | 8626 | 4410 | 4842 | 661 | 7143 | 723 |
| 'Lactobacillales | 'Lactobacillaceae | 'Lactobacillus | '_ | 8555 | 11554 | 3433 | 5804 | 1056 | 2815 | 178 | 3824 | 416 | 254 |
| 'Lactobacillales | 'Streptococcaceae | 'Lactococcus | '_ | 5038 | 1081 | 2755 | 2146 | 4802 | 5395 | 2440 | 2577 | 2239 | 1948 |
| 'Bacteroidales | 'Bacteroidaceae | 'Bacteroides | '_ | 1091 | 1562 | 1707 | 2831 | 2262 | 1319 | 2690 | 2355 | 1596 | 8412 |
| 'Bacteroidales | '[Paraprevotellaceae] | '[Prevotella] | '_ | 962 | 540 | 1777 | 1865 | 1572 | 3708 | 3086 | 1520 | 1674 | 7582 |
| 'Clostridiales | '_ | '_ | '_ | 17 | 5265 | 680 | 5964 | 620 | 837 | 1655 | 988 | 1552 | 6029 |
| 'Deferribacterales | 'Deferribacteraceae | 'Mucispirillum | 'schaedleri | 36 | 2087 | 1486 | 1644 | 205 | 2449 | 2438 | 832 | 4795 | 1921 |
| 'Bacteroidales | 'Bacteroidaceae | 'Bacteroides | 'acidifaciens | 322 | 2633 | 781 | 2161 | 988 | 2498 | 1623 | 2156 | 983 | 2571 |
| 'Clostridiales | '_ | '_ | '_ | 497 | 1386 | 760 | 1478 | 365 | 372 | 2325 | 1398 | 853 | 2764 |
| 'Verrucomicrobiales | 'Verrucomicrobiaceae | 'Akkermansia | 'muciniphila | 2151 | 0 | 1963 | 746 | 186 | 0 | 2422 | 0 | 2305 | 320 |
| 'Bacteroidales | 'Rikenellaceae | '_ | '_ | 475 | 616 | 849 | 769 | 1096 | 2251 | 1147 | 972 | 696 | 1175 |
| 'Enterobacteriales | 'Enterobacteriaceae | 'Other | 'Other | 0 | 100 | 5 | 0 | 43 | 96 | 17 | 9682 | 17 | 40 |
| 'Lactobacillales | 'Lactobacillaceae | 'Lactobacillus | '_ | 1729 | 594 | 1119 | 459 | 1351 | 1409 | 625 | 859 | 883 | 513 |
| 'Bacteroidales | 'S24-7 | '_ | '_ | 592 | 494 | 953 | 796 | 1207 | 1046 | 676 | 1770 | 625 | 411 |
| 'Clostridiales | 'Lachnospiraceae | '_ | '_ | 149 | 826 | 914 | 1194 | 868 | 861 | 553 | 899 | 752 | 1545 |
| 'Clostridiales | 'Peptostreptococcaceae | '_ | '_ | 1705 | 1344 | 7 | 691 | 1190 | 579 | 1287 | 310 | 1121 | 181 |
| 'Clostridiales | 'Lachnospiraceae | '[Ruminococcus] | 'gnavus | 1015 | 582 | 1118 | 398 | 617 | 167 | 1002 | 466 | 2041 | 898 |
| 'Clostridiales | 'Lachnospiraceae | 'Coprococcus | '_ | 90 | 708 | 955 | 1191 | 873 | 917 | 448 | 991 | 444 | 1468 |
| 'Bifidobacteriales | 'Bifidobacteriaceae | 'Bifidobacterium | 'pseudolongum | 711 | 180 | 1236 | 374 | 2092 | 602 | 671 | 82 | 1317 | 111 |
| 'Enterobacteriales | 'Enterobacteriaceae | 'Proteus | '_ | 14 | 1432 | 42 | 520 | 48 | 1123 | 157 | 2722 | 0 | 1153 |
| 'Lactobacillales | 'Lactobacillaceae | 'Lactobacillus | 'Other | 1456 | 2494 | 362 | 844 | 140 | 579 | 55 | 854 | 64 | 111 |
| 'Bacteroidales | 'S24-7 | '_ | '_ | 442 | 819 | 804 | 555 | 993 | 722 | 575 | 784 | 740 | 329 |
| 'Bacteroidales | 'Porphyromonadaceae | 'Parabacteroides | 'distasonis | 111 | 699 | 194 | 700 | 361 | 689 | 330 | 585 | 320 | 1437 |
| 'Clostridiales | 'Clostridiaceae | '_ | '_ | 389 | 387 | 384 | 572 | 1317 | 521 | 384 | 103 | 1242 | 79 |
| 'Clostridiales | '_ | '_ | '_ | 41 | 998 | 289 | 1118 | 98 | 839 | 392 | 421 | 420 | 536 |
| 'Desulfovibrionales | 'Desulfovibrionaceae | 'Desulfovibrio | '_ | 0 | 734 | 8 | 739 | 21 | 1998 | 142 | 850 | 65 | 503 |
| 'Bacteroidales | 'S24-7 | '_ | '_ | 331 | 155 | 629 | 495 | 423 | 1087 | 551 | 327 | 708 | 271 |
| 'Coriobacteriales | 'Coriobacteriaceae | 'Adlercreutzia | '_ | 1261 | 399 | 411 | 280 | 545 | 387 | 456 | 388 | 625 | 135 |
| 'Bacteroidales | 'S24-7 | '_ | '_ | 216 | 471 | 457 | 163 | 831 | 341 | 827 | 530 | 480 | 527 |
| 'Erysipelotrichales | 'Erysipelotrichaceae | 'Clostridium | 'cocleatum | 0 | 519 | 95 | 1185 | 480 | 828 | 52 | 587 | 319 | 696 |
| 'Clostridiales | 'Ruminococcaceae | 'Ruminococcus | '_ | 532 | 50 | 1331 | 257 | 313 | 12 | 650 | 172 | 1164 | 266 |
| 'Clostridiales | '_ | '_ | '_ | 250 | 522 | 302 | 564 | 161 | 144 | 855 | 488 | 319 | 995 |
| 'Clostridiales | 'Lachnospiraceae | 'Other | 'Other | 35 | 611 | 267 | 832 | 332 | 628 | 164 | 606 | 481 | 436 |
| 'Clostridiales | 'Lachnospiraceae | '[Ruminococcus] | 'gnavus | 238 | 0 | 1211 | 25 | 1233 | 0 | 995 | 90 | 311 | 87 |
| 'Turicibacterales | 'Turicibacteraceae | 'Turicibacter | '_ | 822 | 269 | 472 | 206 | 824 | 217 | 627 | 0 | 586 | 0 |
| 'Clostridiales | 'Lachnospiraceae | '_ | '_ | 119 | 119 | 46 | 119 | 721 | 1218 | 638 | 139 | 611 | 253 |
| 'Bacteroidales | 'Bacteroidaceae | 'Bacteroides | '_ | 18 | 180 | 73 | 283 | 316 | 193 | 114 | 190 | 460 | 2042 |
| 'Clostridiales | 'Ruminococcaceae | 'Oscillospira | '_ | 82 | 267 | 400 | 804 | 232 | 162 | 203 | 725 | 293 | 621 |
| 'Clostridiales | 'Clostridiaceae | 'Other | 'Other | 923 | 314 | 362 | 132 | 678 | 388 | 527 | 51 | 317 | 70 |
| 'Clostridiales | 'Lachnospiraceae | 'Roseburia | '_ | 88 | 408 | 466 | 878 | 156 | 255 | 0 | 85 | 488 | 927 |
| 'Clostridiales | 'Lachnospiraceae | '_ | '_ | 66 | 461 | 386 | 658 | 285 | 100 | 349 | 348 | 338 | 698 |
| 'Bacteroidales | 'S24-7 | '_ | '_ | 389 | 98 | 291 | 129 | 556 | 235 | 1359 | 128 | 305 | 104 |
| 'Clostridiales | 'Lachnospiraceae | '[Ruminococcus] | 'gnavus | 44 | 466 | 522 | 0 | 71 | 577 | 216 | 1253 | 123 | 126 |
| 'Clostridiales | '_ | '_ | '_ | 224 | 104 | 692 | 49 | 882 | 13 | 640 | 83 | 418 | 236 |
| 'Bacteroidales | '_ | '_ | '_ | 82 | 323 | 358 | 299 | 407 | 428 | 353 | 404 | 357 | 267 |
| 'Clostridiales | 'Lachnospiraceae | 'Coprococcus | '_ | 38 | 297 | 349 | 454 | 350 | 387 | 122 | 399 | 126 | 562 |
| 'Bacteroidales | 'S24-7 | '_ | '_ | 146 | 199 | 230 | 549 | 215 | 429 | 273 | 567 | 322 | 144 |
| 'Bacteroidales | 'S24-7 | '_ | '_ | 153 | 340 | 338 | 131 | 411 | 262 | 379 | 297 | 237 | 410 |
| 'Clostridiales | '_ | '_ | '_ | 7 | 283 | 199 | 275 | 698 | 342 | 113 | 322 | 116 | 590 |
| 'Clostridiales | 'Lachnospiraceae | 'Other | 'Other | 263 | 169 | 65 | 100 | 511 | 627 | 171 | 218 | 562 | 245 |
| 'Clostridiales | 'Lachnospiraceae | '[Ruminococcus] | 'gnavus | 386 | 62 | 162 | 433 | 227 | 221 | 217 | 119 | 375 | 686 |
| 'Erysipelotrichales | 'Erysipelotrichaceae | 'Allobaculum | '_ | 512 | 501 | 415 | 301 | 199 | 402 | 136 | 102 | 286 | 0 |
| 'Desulfovibrionales | 'Desulfovibrionaceae | 'Desulfovibrio | 'C21_c20 | 47 | 98 | 333 | 294 | 111 | 846 | 307 | 171 | 604 | 43 |
| 'Bacteroidales | 'S24-7 | '_ | '_ | 212 | 120 | 366 | 271 | 198 | 585 | 297 | 255 | 383 | 152 |
| 'Bacteroidales | '_ | '_ | '_ | 53 | 177 | 228 | 178 | 215 | 230 | 380 | 424 | 524 | 345 |
| 'Lactobacillales | 'Streptococcaceae | 'Lactococcus | 'garvieae | 167 | 0 | 0 | 0 | 21 | 24 | 17 | 2375 | 11 | 8 |
| 'Clostridiales | '_ | '_ | '_ | 0 | 0 | 227 | 0 | 2184 | 0 | 0 | 196 | 0 | 0 |
| 'Enterobacteriales | 'Enterobacteriaceae | 'Proteus | '_ | 0 | 507 | 0 | 184 | 0 | 408 | 54 | 1019 | 0 | 386 |
| 'Clostridiales | 'Lachnospiraceae | '_ | '_ | 76 | 103 | 760 | 0 | 387 | 303 | 55 | 25 | 44 | 786 |
| 'Bacteroidales | 'S24-7 | '_ | '_ | 162 | 120 | 376 | 103 | 300 | 188 | 533 | 264 | 399 | 74 |
| 'Clostridiales | 'Lachnospiraceae | '[Ruminococcus] | 'gnavus | 38 | 31 | 471 | 179 | 119 | 802 | 87 | 159 | 278 | 331 |
| 'Clostridiales | '_ | '_ | '_ | 273 | 0 | 690 | 38 | 459 | 53 | 522 | 18 | 371 | 44 |
| 'Clostridiales | 'Clostridiaceae | 'Clostridium | 'Other | 145 | 196 | 179 | 284 | 529 | 258 | 138 | 0 | 707 | 0 |
| 'Clostridiales | 'Lachnospiraceae | '_ | '_ | 37 | 179 | 867 | 356 | 82 | 98 | 774 | 16 | 0 | 15 |
| 'Clostridiales | 'Clostridiaceae | 'Other | 'Other | 542 | 233 | 252 | 97 | 379 | 294 | 356 | 0 | 215 | 49 |
| 'Clostridiales | '_ | '_ | '_ | 403 | 21 | 25 | 0 | 135 | 90 | 49 | 1299 | 313 | 54 |
| 'Clostridiales | 'Clostridiaceae | 'Other | 'Other | 726 | 281 | 307 | 127 | 276 | 149 | 209 | 0 | 255 | 57 |
| 'Bacteroidales | 'S24-7 | '_ | '_ | 163 | 205 | 408 | 199 | 396 | 224 | 209 | 282 | 196 | 101 |
| 'Clostridiales | 'Lachnospiraceae | '[Ruminococcus] | 'gnavus | 301 | 151 | 295 | 118 | 175 | 41 | 296 | 137 | 601 | 260 |
| 'Bacteroidales | 'Bacteroidaceae | 'Bacteroides | '_ | 11 | 20 | 74 | 66 | 40 | 20 | 253 | 94 | 170 | 1596 |
| 'Bacteroidales | 'S24-7 | '_ | '_ | 165 | 67 | 318 | 220 | 210 | 417 | 293 | 193 | 327 | 125 |
| 'Clostridiales | '_ | '_ | '_ | 264 | 0 | 754 | 39 | 401 | 46 | 443 | 10 | 323 | 36 |
| 'Clostridiales | 'Lachnospiraceae | '[Ruminococcus] | 'gnavus | 310 | 177 | 363 | 58 | 305 | 88 | 241 | 323 | 120 | 281 |
| 'Clostridiales | 'Ruminococcaceae | 'Oscillospira | '_ | 69 | 164 | 329 | 288 | 266 | 60 | 291 | 110 | 284 | 393 |
| 'Clostridiales | 'Ruminococcaceae | 'Anaerotruncus | '_ | 38 | 112 | 189 | 113 | 198 | 62 | 429 | 268 | 638 | 122 |
| 'Burkholderiales | 'Alcaligenaceae | 'Sutterella | '_ | 257 | 399 | 217 | 107 | 159 | 148 | 401 | 226 | 191 | 63 |
| 'Bacteroidales | 'S24-7 | '_ | '_ | 169 | 83 | 368 | 85 | 188 | 133 | 439 | 222 | 394 | 54 |
| 'CW040 | 'F16 | '_ | '_ | 510 | 117 | 0 | 0 | 0 | 253 | 161 | 532 | 329 | 229 |
| 'Clostridiales | '_ | '_ | '_ | 305 | 0 | 37 | 0 | 1318 | 0 | 29 | 121 | 320 | 0 |
| 'Clostridiales | 'Other | 'Other | 'Other | 456 | 360 | 0 | 170 | 312 | 151 | 369 | 0 | 276 | 0 |
| 'Clostridiales | 'Lachnospiraceae | '[Ruminococcus] | 'gnavus | 146 | 173 | 175 | 91 | 385 | 316 | 100 | 203 | 168 | 325 |
| 'Bacteroidales | 'S24-7 | '_ | '_ | 206 | 0 | 230 | 0 | 356 | 110 | 427 | 78 | 476 | 153 |
| 'Bacteroidales | 'Rikenellaceae | '_ | '_ | 0 | 135 | 162 | 264 | 181 | 193 | 144 | 460 | 259 | 134 |
| 'Clostridiales | 'Lachnospiraceae | '[Ruminococcus] | 'gnavus | 656 | 492 | 0 | 0 | 0 | 188 | 486 | 16 | 62 | 30 |
| 'Clostridiales | 'Lachnospiraceae | '_ | '_ | 0 | 126 | 118 | 244 | 75 | 266 | 77 | 398 | 250 | 359 |
| 'Clostridiales | 'Clostridiaceae | 'Other | 'Other | 0 | 119 | 164 | 235 | 341 | 270 | 0 | 0 | 693 | 39 |
| 'Clostridiales | 'Lachnospiraceae | '_ | '_ | 46 | 160 | 111 | 205 | 70 | 503 | 87 | 350 | 78 | 239 |
| 'Clostridiales | 'Ruminococcaceae | 'Oscillospira | '_ | 117 | 249 | 97 | 220 | 162 | 264 | 171 | 258 | 160 | 138 |
| 'Lactobacillales | 'Enterococcaceae | 'Enterococcus | '_ | 45 | 822 | 42 | 61 | 183 | 33 | 28 | 486 | 32 | 0 |
| 'Bacteroidales | 'Rikenellaceae | '_ | '_ | 39 | 119 | 147 | 230 | 165 | 162 | 126 | 388 | 239 | 96 |
| 'Bacteroidales | 'S24-7 | '_ | '_ | 188 | 40 | 153 | 59 | 351 | 132 | 401 | 81 | 208 | 79 |
| 'Lactobacillales | 'Lactobacillaceae | 'Lactobacillus | 'reuteri | 517 | 428 | 157 | 340 | 92 | 0 | 0 | 81 | 0 | 0 |
| 'Bacteroidales | 'Bacteroidaceae | 'Bacteroides | 'acidifaciens | 0 | 124 | 90 | 202 | 141 | 132 | 251 | 386 | 161 | 127 |
| 'Clostridiales | 'Ruminococcaceae | 'Oscillospira | '_ | 75 | 51 | 162 | 204 | 98 | 14 | 83 | 237 | 202 | 451 |
| 'Enterobacteriales | 'Enterobacteriaceae | 'Escherichia | 'coli | 0 | 0 | 0 | 0 | 0 | 0 | 0 | 1574 | 0 | 0 |
| 'Clostridiales | 'Ruminococcaceae | 'Oscillospira | '_ | 16 | 135 | 262 | 137 | 31 | 67 | 185 | 308 | 152 | 280 |
| 'Clostridiales | '_ | '_ | '_ | 117 | 29 | 236 | 17 | 288 | 12 | 312 | 42 | 228 | 254 |
| 'Bacteroidales | 'S24-7 | '_ | '_ | 84 | 250 | 173 | 62 | 166 | 126 | 247 | 130 | 157 | 92 |
| 'Clostridiales | '_ | '_ | '_ | 159 | 0 | 0 | 0 | 283 | 32 | 0 | 0 | 988 | 0 |
| 'Clostridiales | 'Other | 'Other | 'Other | 11 | 230 | 149 | 27 | 98 | 39 | 57 | 109 | 463 | 251 |
| 'Clostridiales | 'Lachnospiraceae | '_ | '_ | 51 | 287 | 67 | 224 | 40 | 270 | 47 | 238 | 46 | 163 |
| 'Clostridiales | 'Lachnospiraceae | '[Ruminococcus] | 'gnavus | 88 | 190 | 185 | 272 | 278 | 37 | 79 | 43 | 148 | 63 |
| 'Clostridiales | 'Lachnospiraceae | '_ | '_ | 28 | 17 | 54 | 62 | 51 | 288 | 167 | 158 | 390 | 117 |
| 'Enterobacteriales | 'Enterobacteriaceae | 'Klebsiella | 'Other | 0 | 225 | 0 | 0 | 0 | 0 | 18 | 1056 | 0 | 33 |
| 'Clostridiales | 'Lachnospiraceae | '_ | '_ | 11 | 51 | 105 | 99 | 77 | 364 | 105 | 229 | 76 | 187 |
| 'Enterobacteriales | 'Enterobacteriaceae | 'Klebsiella | '_ | 0 | 210 | 0 | 0 | 0 | 0 | 15 | 1051 | 0 | 27 |
| 'Clostridiales | '_ | '_ | '_ | 11 | 32 | 491 | 61 | 37 | 226 | 48 | 128 | 0 | 261 |
| 'Lactobacillales | 'Lactobacillaceae | 'Lactobacillus | '_ | 167 | 857 | 26 | 183 | 21 | 0 | 0 | 0 | 4 | 0 |
| 'Clostridiales | 'Other | 'Other | 'Other | 27 | 102 | 233 | 117 | 106 | 50 | 129 | 183 | 57 | 248 |
| 'Clostridiales | '_ | '_ | '_ | 0 | 176 | 114 | 243 | 100 | 0 | 191 | 213 | 209 | 0 |
| 'Erysipelotrichales | 'Erysipelotrichaceae | 'Allobaculum | '_ | 0 | 0 | 0 | 0 | 6 | 978 | 0 | 0 | 203 | 59 |
| 'Clostridiales | '_ | '_ | '_ | 72 | 0 | 267 | 0 | 368 | 0 | 264 | 49 | 131 | 84 |
| 'Clostridiales | 'Lachnospiraceae | '_ | '_ | 34 | 15 | 188 | 165 | 115 | 48 | 161 | 19 | 207 | 271 |
| 'Clostridiales | '_ | '_ | '_ | 4 | 263 | 54 | 8 | 0 | 0 | 107 | 246 | 362 | 173 |
| 'Clostridiales | 'Other | 'Other | 'Other | 18 | 149 | 70 | 183 | 42 | 179 | 71 | 168 | 58 | 269 |
| 'Lactobacillales | 'Lactobacillaceae | 'Lactobacillus | 'reuteri | 339 | 470 | 55 | 93 | 0 | 92 | 0 | 157 | 0 | 0 |
| 'Clostridiales | 'Ruminococcaceae | 'Oscillospira | '_ | 40 | 77 | 88 | 244 | 78 | 0 | 123 | 123 | 158 | 273 |
| 'Clostridiales | 'Lachnospiraceae | '_ | '_ | 27 | 122 | 93 | 193 | 74 | 147 | 142 | 211 | 50 | 139 |
| 'Bacteroidales | 'S24-7 | '_ | '_ | 90 | 0 | 0 | 103 | 0 | 254 | 134 | 318 | 158 | 125 |
| 'Bacteroidales | 'S24-7 | '_ | '_ | 91 | 55 | 97 | 87 | 190 | 87 | 150 | 154 | 64 | 111 |
| 'Clostridiales | 'Lachnospiraceae | '_ | '_ | 22 | 31 | 69 | 45 | 109 | 64 | 257 | 54 | 315 | 119 |
| 'Bacteroidales | 'S24-7 | '_ | '_ | 40 | 60 | 75 | 198 | 84 | 158 | 90 | 216 | 104 | 58 |
| 'Bacteroidales | 'S24-7 | '_ | '_ | 123 | 36 | 161 | 39 | 95 | 261 | 151 | 82 | 60 | 74 |
| 'Clostridiales | 'Lachnospiraceae | '_ | '_ | 22 | 229 | 165 | 192 | 76 | 143 | 66 | 66 | 19 | 98 |
| 'Clostridiales | '_ | '_ | '_ | 22 | 40 | 18 | 109 | 12 | 39 | 50 | 370 | 28 | 376 |
| 'Coriobacteriales | 'Coriobacteriaceae | 'Adlercreutzia | '_ | 121 | 63 | 196 | 33 | 270 | 21 | 115 | 98 | 110 | 30 |
| 'Clostridiales | '_ | '_ | '_ | 43 | 53 | 83 | 95 | 287 | 0 | 74 | 101 | 200 | 118 |
| 'Clostridiales | 'Clostridiaceae | 'Other | 'Other | 100 | 0 | 0 | 0 | 362 | 211 | 330 | 47 | 0 | 0 |
| 'Turicibacterales | 'Turicibacteraceae | 'Turicibacter | '_ | 521 | 0 | 0 | 0 | 230 | 0 | 140 | 0 | 125 | 0 |
| 'Clostridiales | 'Ruminococcaceae | 'Ruminococcus | '_ | 49 | 0 | 105 | 81 | 199 | 47 | 117 | 64 | 193 | 143 |
| 'Clostridiales | 'Ruminococcaceae | 'Oscillospira | '_ | 13 | 36 | 105 | 75 | 193 | 13 | 152 | 67 | 215 | 119 |
| 'Clostridiales | 'Dehalobacteriaceae | 'Dehalobacterium | '_ | 20 | 100 | 38 | 256 | 0 | 118 | 29 | 94 | 43 | 268 |
| 'Bacillales | 'Staphylococcaceae | 'Staphylococcus | 'Other | 177 | 88 | 72 | 81 | 156 | 109 | 35 | 89 | 110 | 34 |
| 'Bacteroidales | 'Rikenellaceae | '_ | '_ | 13 | 139 | 62 | 140 | 55 | 182 | 64 | 121 | 61 | 97 |
| 'Clostridiales | 'Lachnospiraceae | '_ | '_ | 155 | 63 | 109 | 114 | 109 | 63 | 85 | 43 | 87 | 98 |
| 'Clostridiales | '_ | '_ | '_ | 53 | 0 | 188 | 0 | 289 | 0 | 261 | 0 | 60 | 68 |
| 'Bacillales | 'Staphylococcaceae | 'Staphylococcus | 'sciuri | 184 | 139 | 66 | 63 | 124 | 63 | 131 | 54 | 34 | 41 |
| 'Clostridiales | '_ | '_ | '_ | 356 | 25 | 0 | 0 | 346 | 18 | 0 | 40 | 103 | 0 |
| 'Clostridiales | 'Ruminococcaceae | 'Ruminococcus | '_ | 72 | 16 | 98 | 119 | 120 | 4 | 117 | 61 | 157 | 124 |
| 'Coriobacteriales | 'Coriobacteriaceae | 'Adlercreutzia | '_ | 176 | 34 | 96 | 46 | 190 | 70 | 77 | 84 | 101 | 0 |
| 'Clostridiales | 'Ruminococcaceae | 'Oscillospira | '_ | 34 | 103 | 67 | 101 | 92 | 117 | 91 | 80 | 61 | 120 |
| 'Clostridiales | 'Lachnospiraceae | 'Clostridium | 'Other | 0 | 15 | 32 | 15 | 109 | 387 | 102 | 72 | 85 | 40 |
| 'Clostridiales | 'Lachnospiraceae | 'Coprococcus | '_ | 56 | 69 | 92 | 61 | 65 | 58 | 67 | 145 | 151 | 85 |
| 'Desulfovibrionales | 'Desulfovibrionaceae | '_ | '_ | 4 | 90 | 41 | 188 | 22 | 171 | 41 | 94 | 50 | 148 |
| 'Bacteroidales | '_ | '_ | '_ | 32 | 28 | 162 | 23 | 149 | 61 | 92 | 104 | 94 | 103 |
| 'Clostridiales | 'Lachnospiraceae | '_ | '_ | 22 | 53 | 255 | 0 | 142 | 99 | 0 | 0 | 0 | 272 |
| 'Clostridiales | 'Dehalobacteriaceae | 'Dehalobacterium | '_ | 0 | 98 | 34 | 203 | 33 | 80 | 24 | 86 | 41 | 239 |
| 'Coriobacteriales | 'Coriobacteriaceae | 'Adlercreutzia | '_ | 108 | 52 | 127 | 63 | 214 | 20 | 79 | 83 | 90 | 0 |
| 'Clostridiales | 'Lachnospiraceae | '_ | '_ | 14 | 63 | 39 | 108 | 22 | 204 | 42 | 168 | 37 | 119 |
| 'Clostridiales | 'Lachnospiraceae | '_ | '_ | 87 | 48 | 193 | 17 | 82 | 37 | 97 | 70 | 168 | 14 |
| 'Clostridiales | 'Ruminococcaceae | 'Oscillospira | '_ | 0 | 0 | 0 | 101 | 0 | 0 | 0 | 220 | 0 | 453 |
| 'Bacteroidales | 'S24-7 | '_ | '_ | 56 | 33 | 158 | 47 | 129 | 173 | 57 | 0 | 55 | 51 |
| 'Lactobacillales | 'Streptococcaceae | 'Streptococcus | '_ | 0 | 112 | 0 | 106 | 0 | 320 | 0 | 165 | 0 | 44 |
| 'Clostridiales | 'Lachnospiraceae | '[Ruminococcus] | 'gnavus | 0 | 126 | 107 | 165 | 171 | 0 | 51 | 0 | 83 | 37 |
| 'Clostridiales | 'Lachnospiraceae | '_ | '_ | 0 | 0 | 144 | 102 | 60 | 18 | 101 | 17 | 102 | 173 |
| 'Clostridiales | 'Lachnospiraceae | '_ | '_ | 50 | 11 | 158 | 17 | 88 | 11 | 133 | 4 | 231 | 10 |
| 'Coriobacteriales | 'Coriobacteriaceae | 'Adlercreutzia | '_ | 142 | 26 | 59 | 19 | 160 | 49 | 84 | 76 | 74 | 8 |
| 'Clostridiales | 'Lachnospiraceae | '_ | '_ | 220 | 55 | 7 | 52 | 22 | 135 | 17 | 94 | 25 | 57 |
| 'Erysipelotrichales | 'Erysipelotrichaceae | 'Clostridium | 'cocleatum | 0 | 106 | 0 | 222 | 74 | 130 | 0 | 0 | 0 | 119 |
| 'Bacteroidales | 'S24-7 | '_ | '_ | 15 | 93 | 36 | 160 | 31 | 85 | 36 | 86 | 53 | 50 |
| 'Clostridiales | 'Lachnospiraceae | '_ | '_ | 7 | 64 | 40 | 61 | 18 | 156 | 38 | 142 | 27 | 81 |
| 'Bacteroidales | 'Bacteroidaceae | 'Bacteroides | '_ | 0 | 0 | 0 | 0 | 0 | 0 | 77 | 31 | 55 | 468 |
| 'Clostridiales | 'Ruminococcaceae | 'Oscillospira | '_ | 0 | 87 | 46 | 97 | 65 | 45 | 60 | 110 | 59 | 53 |
| 'Clostridiales | 'Ruminococcaceae | '_ | '_ | 21 | 81 | 41 | 139 | 27 | 91 | 38 | 67 | 41 | 72 |
| 'Clostridiales | 'Lachnospiraceae | 'Other | 'Other | 42 | 53 | 11 | 86 | 46 | 82 | 39 | 114 | 52 | 90 |
| 'Clostridiales | 'Lachnospiraceae | 'Dorea | '_ | 23 | 50 | 46 | 63 | 31 | 22 | 67 | 129 | 28 | 154 |
| 'Clostridiales | 'Lachnospiraceae | '_ | '_ | 0 | 70 | 0 | 78 | 0 | 203 | 0 | 168 | 0 | 94 |
| 'Bacteroidales | 'S24-7 | '_ | '_ | 34 | 0 | 130 | 29 | 120 | 0 | 74 | 42 | 168 | 0 |
| 'Clostridiales | 'Lachnospiraceae | '_ | '_ | 0 | 0 | 87 | 96 | 45 | 24 | 89 | 14 | 92 | 138 |
| 'Lactobacillales | 'Lactobacillaceae | 'Lactobacillus | 'Other | 107 | 83 | 148 | 189 | 52 | 0 | 0 | 0 | 0 | 0 |
| 'Clostridiales | '_ | '_ | '_ | 0 | 29 | 39 | 73 | 23 | 85 | 32 | 212 | 34 | 48 |
| 'Clostridiales | 'Ruminococcaceae | 'Oscillospira | '_ | 24 | 0 | 95 | 52 | 80 | 0 | 108 | 0 | 99 | 101 |
| 'Bacteroidales | 'S24-7 | '_ | '_ | 0 | 64 | 43 | 52 | 65 | 92 | 53 | 52 | 55 | 79 |
| 'Bacteroidales | 'S24-7 | '_ | '_ | 55 | 0 | 52 | 53 | 76 | 70 | 54 | 53 | 67 | 71 |
| 'Clostridiales | '[Mogibacteriaceae] | '_ | '_ | 55 | 44 | 76 | 53 | 100 | 64 | 22 | 64 | 36 | 28 |
| 'Clostridiales | 'Peptococcaceae | '_ | '_ | 18 | 36 | 80 | 152 | 58 | 8 | 74 | 16 | 43 | 42 |
| 'Clostridiales | '_ | '_ | '_ | 42 | 0 | 42 | 0 | 312 | 8 | 43 | 23 | 13 | 40 |
| 'Clostridiales | '_ | '_ | '_ | 0 | 83 | 36 | 0 | 48 | 0 | 65 | 72 | 0 | 213 |
| 'Bacteroidales | 'S24-7 | '_ | '_ | 31 | 33 | 99 | 36 | 100 | 0 | 76 | 45 | 76 | 19 |
| 'Clostridiales | 'Ruminococcaceae | 'Oscillospira | '_ | 17 | 26 | 25 | 88 | 39 | 6 | 44 | 49 | 85 | 135 |
| 'Clostridiales | 'Ruminococcaceae | 'Oscillospira | '_ | 15 | 4 | 110 | 14 | 95 | 0 | 76 | 45 | 97 | 53 |
| 'Clostridiales | 'Lachnospiraceae | '[Ruminococcus] | 'gnavus | 43 | 0 | 135 | 0 | 81 | 0 | 82 | 39 | 33 | 87 |
| 'Clostridiales | 'Ruminococcaceae | 'Oscillospira | '_ | 15 | 33 | 35 | 100 | 53 | 93 | 26 | 35 | 57 | 42 |
| 'Lactobacillales | 'Lactobacillaceae | 'Lactobacillus | 'reuteri | 80 | 76 | 103 | 178 | 41 | 0 | 0 | 0 | 0 | 0 |
| 'Coriobacteriales | 'Coriobacteriaceae | 'Adlercreutzia | '_ | 112 | 0 | 73 | 0 | 105 | 57 | 71 | 0 | 58 | 0 |
| 'Clostridiales | 'Lachnospiraceae | '_ | '_ | 12 | 36 | 0 | 0 | 0 | 0 | 66 | 362 | 0 | 0 |
| 'Bacteroidales | 'S24-7 | '_ | '_ | 0 | 48 | 0 | 113 | 48 | 152 | 0 | 0 | 64 | 49 |
| 'Turicibacterales | 'Turicibacteraceae | 'Turicibacter | '_ | 0 | 134 | 0 | 0 | 0 | 0 | 339 | 0 | 0 | 0 |
| 'Bacteroidales | 'S24-7 | '_ | '_ | 0 | 128 | 0 | 53 | 62 | 139 | 0 | 0 | 45 | 41 |
| 'Bacteroidales | 'Rikenellaceae | '_ | '_ | 11 | 84 | 14 | 84 | 23 | 50 | 22 | 61 | 25 | 86 |
| 'Enterobacteriales | 'Enterobacteriaceae | 'Other | 'Other | 0 | 99 | 0 | 0 | 0 | 0 | 0 | 357 | 0 | 0 |
| 'Bacteroidales | 'Rikenellaceae | 'Other | 'Other | 5 | 37 | 40 | 43 | 38 | 102 | 16 | 72 | 47 | 53 |
| 'Bacteroidales | 'Porphyromonadaceae | 'Parabacteroides | '_ | 0 | 63 | 25 | 72 | 18 | 59 | 25 | 81 | 16 | 79 |
| 'Lactobacillales | 'Lactobacillaceae | 'Lactobacillus | 'Other | 87 | 81 | 86 | 146 | 36 | 0 | 0 | 0 | 0 | 0 |
| 'Clostridiales | '_ | '_ | '_ | 0 | 0 | 47 | 38 | 213 | 15 | 27 | 34 | 60 | 0 |
| 'Bacteroidales | 'S24-7 | '_ | '_ | 13 | 22 | 53 | 73 | 29 | 45 | 47 | 78 | 27 | 45 |
| 'Bacteroidales | 'S24-7 | '_ | '_ | 15 | 0 | 30 | 83 | 33 | 49 | 23 | 53 | 35 | 109 |
| 'Clostridiales | '_ | '_ | '_ | 8 | 43 | 50 | 14 | 40 | 32 | 18 | 121 | 85 | 19 |
| 'Enterobacteriales | 'Enterobacteriaceae | 'Klebsiella | 'Other | 0 | 82 | 0 | 0 | 0 | 0 | 0 | 348 | 0 | 0 |
| 'Clostridiales | '_ | '_ | '_ | 32 | 0 | 73 | 5 | 112 | 12 | 96 | 31 | 63 | 3 |
| 'Clostridiales | 'Lachnospiraceae | 'Other | 'Other | 8 | 24 | 65 | 67 | 32 | 0 | 18 | 23 | 48 | 137 |
| 'Clostridiales | 'Ruminococcaceae | 'Oscillospira | '_ | 0 | 102 | 0 | 134 | 0 | 0 | 56 | 120 | 0 | 0 |
| 'Clostridiales | 'Ruminococcaceae | 'Oscillospira | '_ | 0 | 0 | 67 | 30 | 54 | 66 | 54 | 0 | 71 | 65 |
| 'Bacteroidales | 'Bacteroidaceae | 'Bacteroides | 'acidifaciens | 0 | 31 | 0 | 35 | 0 | 47 | 76 | 145 | 27 | 35 |
| 'Erysipelotrichales | 'Erysipelotrichaceae | '_ | '_ | 85 | 7 | 29 | 14 | 25 | 114 | 31 | 39 | 51 | 0 |
| 'Clostridiales | 'Lachnospiraceae | 'Dorea | '_ | 18 | 6 | 0 | 15 | 92 | 44 | 18 | 30 | 98 | 74 |
| 'Enterobacteriales | 'Enterobacteriaceae | 'Enterobacter | 'Other | 125 | 30 | 0 | 7 | 0 | 49 | 17 | 125 | 0 | 41 |
| 'Clostridiales | 'Ruminococcaceae | 'Oscillospira | '_ | 0 | 0 | 0 | 82 | 0 | 0 | 0 | 225 | 42 | 45 |
| 'Clostridiales | 'Ruminococcaceae | 'Other | 'Other | 12 | 60 | 63 | 44 | 60 | 27 | 43 | 10 | 61 | 12 |
| 'Clostridiales | 'Ruminococcaceae | 'Oscillospira | '_ | 0 | 0 | 0 | 68 | 0 | 0 | 0 | 134 | 0 | 184 |
| 'Bacteroidales | 'Rikenellaceae | '_ | '_ | 0 | 61 | 0 | 55 | 32 | 110 | 0 | 48 | 31 | 43 |
| 'Clostridiales | '_ | '_ | '_ | 0 | 12 | 63 | 45 | 0 | 8 | 59 | 47 | 59 | 79 |
| 'Clostridiales | 'Lachnospiraceae | 'Other | 'Other | 0 | 0 | 0 | 0 | 0 | 0 | 0 | 0 | 98 | 273 |
| 'Clostridiales | 'Other | 'Other | 'Other | 0 | 39 | 0 | 65 | 0 | 92 | 0 | 60 | 18 | 87 |
| 'Clostridiales | 'Ruminococcaceae | 'Oscillospira | '_ | 0 | 0 | 47 | 37 | 0 | 0 | 94 | 31 | 79 | 72 |
| 'Bacteroidales | 'S24-7 | '_ | '_ | 0 | 28 | 23 | 78 | 34 | 21 | 0 | 46 | 38 | 85 |
| 'Clostridiales | 'Lachnospiraceae | '_ | '_ | 6 | 111 | 15 | 127 | 5 | 0 | 11 | 65 | 3 | 0 |
| 'Bacillales | 'Staphylococcaceae | 'Staphylococcus | '_ | 53 | 168 | 0 | 30 | 0 | 59 | 0 | 0 | 0 | 27 |
| 'Clostridiales | '_ | '_ | '_ | 0 | 0 | 77 | 0 | 87 | 0 | 66 | 0 | 96 | 6 |
| 'Clostridiales | '_ | '_ | '_ | 0 | 60 | 0 | 0 | 16 | 0 | 83 | 81 | 87 | 0 |
| 'Clostridiales | '_ | '_ | '_ | 0 | 0 | 0 | 219 | 0 | 0 | 0 | 0 | 0 | 101 |
| 'Clostridiales | 'Other | 'Other | 'Other | 0 | 15 | 50 | 29 | 33 | 0 | 24 | 77 | 10 | 71 |
| 'Lactobacillales | 'Aerococcaceae | 'Aerococcus | '_ | 7 | 22 | 0 | 85 | 0 | 4 | 0 | 0 | 0 | 187 |
| 'Clostridiales | 'Ruminococcaceae | '_ | '_ | 0 | 0 | 0 | 0 | 264 | 0 | 33 | 0 | 0 | 0 |
| 'Clostridiales | '_ | '_ | '_ | 0 | 0 | 0 | 111 | 0 | 17 | 0 | 0 | 61 | 108 |
| 'Bacillales | 'Staphylococcaceae | 'Staphylococcus | '_ | 40 | 120 | 18 | 0 | 0 | 0 | 28 | 39 | 0 | 36 |
| 'Clostridiales | 'Ruminococcaceae | 'Oscillospira | '_ | 19 | 14 | 31 | 71 | 25 | 0 | 55 | 36 | 0 | 26 |
| 'Clostridiales | '_ | '_ | '_ | 11 | 7 | 36 | 11 | 46 | 0 | 89 | 24 | 0 | 51 |
| 'Clostridiales | 'Ruminococcaceae | 'Ruminococcus | '_ | 6 | 13 | 15 | 110 | 0 | 12 | 11 | 63 | 0 | 43 |
| 'RF39 | '_ | '_ | '_ | 0 | 0 | 0 | 0 | 0 | 0 | 0 | 0 | 6 | 266 |
| 'Clostridiales | 'Other | 'Other | 'Other | 0 | 0 | 19 | 64 | 0 | 140 | 0 | 0 | 0 | 48 |
| 'Clostridiales | 'Ruminococcaceae | 'Oscillospira | '_ | 12 | 0 | 86 | 63 | 39 | 0 | 15 | 25 | 30 | 0 |
| 'Clostridiales | 'Lachnospiraceae | '[Ruminococcus] | 'gnavus | 51 | 18 | 0 | 26 | 0 | 0 | 22 | 26 | 27 | 95 |
| 'Clostridiales | 'Lachnospiraceae | '_ | '_ | 0 | 65 | 0 | 48 | 0 | 101 | 0 | 46 | 0 | 0 |
| 'Clostridiales | 'Other | 'Other | 'Other | 0 | 0 | 0 | 81 | 0 | 134 | 0 | 0 | 0 | 45 |
| 'Clostridiales | 'Ruminococcaceae | 'Oscillospira | '_ | 11 | 23 | 25 | 16 | 56 | 18 | 32 | 18 | 37 | 23 |
| 'Clostridiales | '_ | '_ | '_ | 0 | 0 | 0 | 99 | 0 | 0 | 0 | 0 | 69 | 90 |
| 'Clostridiales | 'Other | 'Other | 'Other | 38 | 46 | 0 | 67 | 0 | 41 | 21 | 17 | 0 | 11 |
| 'Clostridiales | 'Other | 'Other | 'Other | 0 | 0 | 0 | 56 | 0 | 131 | 0 | 0 | 0 | 53 |
| 'Clostridiales | 'Other | 'Other | 'Other | 0 | 0 | 40 | 0 | 0 | 36 | 35 | 0 | 68 | 57 |
| 'Clostridiales | 'Lachnospiraceae | 'Clostridium | 'Other | 0 | 18 | 0 | 0 | 0 | 78 | 0 | 38 | 0 | 101 |
| 'Clostridiales | 'Lachnospiraceae | '_ | '_ | 0 | 16 | 0 | 0 | 12 | 18 | 0 | 164 | 0 | 23 |
| 'Clostridiales | 'Other | 'Other | 'Other | 0 | 18 | 42 | 62 | 8 | 50 | 11 | 27 | 13 | 0 |
| 'Clostridiales | 'Other | 'Other | 'Other | 9 | 40 | 0 | 0 | 0 | 0 | 33 | 71 | 23 | 51 |
| 'Clostridiales | 'Ruminococcaceae | 'Oscillospira | '_ | 0 | 0 | 20 | 62 | 14 | 0 | 17 | 50 | 25 | 38 |
| 'Desulfovibrionales | 'Desulfovibrionaceae | '_ | '_ | 0 | 0 | 0 | 54 | 0 | 62 | 0 | 44 | 23 | 42 |
| 'Clostridiales | 'Dehalobacteriaceae | 'Dehalobacterium | '_ | 11 | 0 | 31 | 0 | 97 | 0 | 56 | 0 | 24 | 0 |
| 'Bacteroidales | 'Porphyromonadaceae | 'Parabacteroides | '_ | 0 | 29 | 9 | 19 | 0 | 17 | 0 | 24 | 3 | 115 |
| 'Clostridiales | 'Ruminococcaceae | 'Ruminococcus | '_ | 0 | 15 | 21 | 38 | 21 | 0 | 26 | 28 | 44 | 22 |
| 'Clostridiales | 'Lachnospiraceae | 'Coprococcus | '_ | 10 | 6 | 39 | 12 | 9 | 12 | 34 | 21 | 23 | 46 |
| 'Clostridiales | 'Ruminococcaceae | 'Oscillospira | '_ | 0 | 0 | 8 | 0 | 0 | 0 | 8 | 119 | 0 | 77 |
| 'Clostridiales | 'Other | 'Other | 'Other | 0 | 0 | 0 | 70 | 0 | 102 | 0 | 0 | 0 | 38 |
| 'Clostridiales | 'Lachnospiraceae | 'Coprococcus | '_ | 0 | 25 | 16 | 0 | 14 | 0 | 19 | 46 | 34 | 49 |
| 'Clostridiales | 'Ruminococcaceae | 'Oscillospira | '_ | 0 | 0 | 0 | 67 | 43 | 0 | 0 | 0 | 87 | 0 |
| 'Clostridiales | 'Christensenellaceae | '_ | '_ | 49 | 8 | 0 | 15 | 3 | 10 | 45 | 19 | 37 | 9 |
| 'Bacteroidales | 'S24-7 | '_ | '_ | 0 | 8 | 11 | 19 | 24 | 0 | 31 | 49 | 44 | 8 |
| 'Clostridiales | 'Ruminococcaceae | 'Oscillospira | '_ | 0 | 0 | 0 | 54 | 43 | 0 | 0 | 0 | 95 | 0 |
| 'Clostridiales | 'Lachnospiraceae | 'Coprococcus | '_ | 0 | 19 | 0 | 54 | 0 | 0 | 0 | 78 | 11 | 29 |
| 'Clostridiales | 'Lachnospiraceae | 'Other | 'Other | 0 | 0 | 0 | 0 | 82 | 102 | 0 | 0 | 0 | 0 |
| 'Lactobacillales | 'Enterococcaceae | 'Vagococcus | '_ | 11 | 0 | 0 | 14 | 0 | 0 | 0 | 138 | 0 | 20 |
| 'Erysipelotrichales | 'Erysipelotrichaceae | 'Clostridium | 'cocleatum | 0 | 28 | 0 | 61 | 0 | 42 | 0 | 0 | 16 | 34 |
| 'Clostridiales | '_ | '_ | '_ | 9 | 0 | 25 | 0 | 29 | 0 | 23 | 0 | 92 | 0 |
| 'Desulfovibrionales | 'Desulfovibrionaceae | 'Bilophila | '_ | 0 | 13 | 8 | 46 | 16 | 0 | 25 | 36 | 11 | 17 |
| 'Clostridiales | '_ | '_ | '_ | 0 | 0 | 22 | 0 | 13 | 0 | 26 | 0 | 107 | 0 |
| 'Actinomycetales | 'Corynebacteriaceae | 'Corynebacterium | 'stationis | 72 | 18 | 0 | 23 | 14 | 0 | 23 | 17 | 0 | 0 |
| 'Clostridiales | 'Lachnospiraceae | 'Other | 'Other | 0 | 41 | 43 | 0 | 0 | 0 | 45 | 0 | 0 | 34 |
| 'Clostridiales | 'Lachnospiraceae | 'Other | 'Other | 0 | 27 | 0 | 26 | 0 | 0 | 0 | 21 | 0 | 82 |
| 'Clostridiales | 'Lachnospiraceae | '_ | '_ | 22 | 7 | 9 | 0 | 17 | 6 | 10 | 59 | 14 | 10 |
| 'Clostridiales | 'Lachnospiraceae | '_ | '_ | 0 | 26 | 6 | 0 | 32 | 0 | 20 | 38 | 31 | 0 |
| 'Clostridiales | '_ | '_ | '_ | 0 | 0 | 22 | 0 | 0 | 0 | 22 | 0 | 108 | 0 |
| 'Clostridiales | '_ | '_ | '_ | 0 | 0 | 0 | 67 | 0 | 0 | 0 | 0 | 0 | 84 |
| 'Clostridiales | '_ | '_ | '_ | 0 | 0 | 30 | 0 | 22 | 0 | 67 | 31 | 0 | 0 |
| 'Clostridiales | 'Ruminococcaceae | '_ | '_ | 5 | 0 | 52 | 0 | 41 | 0 | 23 | 0 | 27 | 0 |
| 'Clostridiales | '_ | '_ | '_ | 0 | 0 | 24 | 0 | 0 | 0 | 25 | 0 | 96 | 0 |
| 'Clostridiales | 'Lachnospiraceae | '_ | '_ | 0 | 0 | 54 | 0 | 0 | 0 | 0 | 0 | 90 | 0 |
| 'Clostridiales | 'Dehalobacteriaceae | 'Dehalobacterium | '_ | 3 | 19 | 8 | 23 | 0 | 26 | 7 | 7 | 3 | 43 |
| 'Clostridiales | 'Other | 'Other | 'Other | 0 | 0 | 71 | 0 | 0 | 0 | 0 | 0 | 0 | 68 |
| 'Lactobacillales | 'Streptococcaceae | 'Streptococcus | 'luteciae | 25 | 0 | 19 | 15 | 27 | 35 | 0 | 5 | 12 | 0 |
| 'Clostridiales | '_ | '_ | '_ | 0 | 0 | 55 | 82 | 0 | 0 | 0 | 0 | 0 | 0 |
| 'Bacteroidales | 'S24-7 | '_ | '_ | 9 | 38 | 0 | 0 | 17 | 0 | 0 | 34 | 0 | 36 |
| 'Clostridiales | 'Ruminococcaceae | 'Anaerotruncus | '_ | 0 | 13 | 7 | 23 | 0 | 25 | 19 | 19 | 6 | 11 |
| 'Clostridiales | 'Ruminococcaceae | 'Ruminococcus | '_ | 0 | 0 | 63 | 0 | 33 | 0 | 16 | 0 | 10 | 0 |
| 'Clostridiales | 'Lachnospiraceae | 'Clostridium | 'Other | 0 | 0 | 0 | 0 | 0 | 96 | 0 | 0 | 26 | 0 |
| 'Clostridiales | '_ | '_ | '_ | 0 | 0 | 31 | 0 | 54 | 0 | 14 | 0 | 10 | 10 |
| 'Clostridiales | 'Ruminococcaceae | 'Oscillospira | '_ | 0 | 6 | 13 | 18 | 19 | 0 | 18 | 11 | 11 | 22 |
| 'Clostridiales | 'Lachnospiraceae | 'Coprococcus | '_ | 0 | 19 | 13 | 25 | 0 | 0 | 11 | 0 | 20 | 30 |
| 'Clostridiales | '_ | '_ | '_ | 20 | 12 | 39 | 0 | 0 | 0 | 32 | 0 | 12 | 0 |
| 'Bacteroidales | 'S24-7 | '_ | '_ | 0 | 0 | 0 | 113 | 0 | 0 | 0 | 0 | 0 | 0 |
| 'Lactobacillales | 'Aerococcaceae | 'Facklamia | '_ | 21 | 0 | 25 | 2 | 27 | 0 | 20 | 0 | 16 | 0 |
| 'Clostridiales | 'Lachnospiraceae | '_ | '_ | 0 | 0 | 0 | 0 | 0 | 0 | 48 | 0 | 0 | 60 |
| 'Clostridiales | 'Ruminococcaceae | 'Oscillospira | '_ | 0 | 0 | 0 | 0 | 0 | 0 | 24 | 0 | 54 | 30 |
| 'Bacteroidales | 'S24-7 | '_ | '_ | 0 | 0 | 0 | 6 | 14 | 6 | 0 | 34 | 47 | 0 |
| 'Clostridiales | '_ | '_ | '_ | 0 | 0 | 0 | 0 | 0 | 0 | 0 | 0 | 0 | 107 |
| 'Clostridiales | 'Ruminococcaceae | 'Oscillospira | '_ | 0 | 0 | 0 | 27 | 5 | 0 | 0 | 29 | 35 | 9 |
| 'Bacteroidales | 'Bacteroidaceae | 'Bacteroides | 'acidifaciens | 0 | 105 | 0 | 0 | 0 | 0 | 0 | 0 | 0 | 0 |
| 'Bacteroidales | 'Bacteroidaceae | 'Bacteroides | 'acidifaciens | 0 | 3 | 6 | 32 | 9 | 17 | 7 | 17 | 0 | 11 |
| 'Clostridiales | 'Lachnospiraceae | 'Dorea | '_ | 11 | 3 | 21 | 0 | 14 | 0 | 19 | 0 | 25 | 8 |
| 'Clostridiales | 'Ruminococcaceae | 'Other | 'Other | 4 | 0 | 14 | 8 | 33 | 0 | 7 | 13 | 10 | 11 |
| 'Clostridiales | 'Lachnospiraceae | '_ | '_ | 0 | 0 | 80 | 0 | 0 | 0 | 0 | 0 | 7 | 13 |
| 'Clostridiales | 'Lachnospiraceae | '_ | '_ | 11 | 0 | 0 | 0 | 30 | 12 | 14 | 12 | 20 | 0 |
| 'Bacteroidales | 'S24-7 | '_ | '_ | 20 | 0 | 0 | 0 | 0 | 3 | 46 | 26 | 1 | 1 |
| 'Clostridiales | 'Lachnospiraceae | 'Other | 'Other | 0 | 0 | 19 | 0 | 0 | 0 | 64 | 0 | 14 | 0 |
| 'Clostridiales | '_ | '_ | '_ | 4 | 0 | 42 | 0 | 50 | 0 | 0 | 0 | 0 | 0 |
| 'Clostridiales | 'Ruminococcaceae | 'Oscillospira | '_ | 0 | 9 | 13 | 15 | 10 | 0 | 0 | 39 | 0 | 9 |
| 'Clostridiales | 'Ruminococcaceae | 'Oscillospira | '_ | 0 | 0 | 0 | 0 | 21 | 0 | 32 | 0 | 42 | 0 |
| 'Clostridiales | 'Lachnospiraceae | '_ | '_ | 0 | 13 | 0 | 20 | 9 | 6 | 13 | 0 | 14 | 19 |
| 'Erysipelotrichales | 'Erysipelotrichaceae | '_ | '_ | 38 | 0 | 0 | 0 | 0 | 55 | 0 | 0 | 0 | 0 |
| 'Bacteroidales | 'S24-7 | '_ | '_ | 0 | 0 | 0 | 0 | 0 | 28 | 0 | 0 | 65 | 0 |
| 'Clostridiales | 'Lachnospiraceae | '_ | '_ | 0 | 0 | 11 | 17 | 18 | 5 | 8 | 11 | 13 | 9 |
| 'Clostridiales | 'Ruminococcaceae | 'Oscillospira | '_ | 15 | 0 | 76 | 0 | 0 | 0 | 0 | 0 | 0 | 0 |
| 'Clostridiales | 'Lachnospiraceae | '_ | '_ | 0 | 0 | 21 | 0 | 8 | 9 | 15 | 0 | 22 | 14 |
| 'Clostridiales | 'Other | 'Other | 'Other | 0 | 0 | 0 | 0 | 0 | 0 | 40 | 0 | 49 | 0 |
| 'CW040 | 'F16 | '_ | '_ | 0 | 0 | 66 | 18 | 0 | 0 | 0 | 0 | 0 | 0 |
| 'Clostridiales | 'Christensenellaceae | '_ | '_ | 6 | 0 | 17 | 5 | 13 | 5 | 9 | 4 | 19 | 5 |
| 'Clostridiales | 'Lachnospiraceae | 'Other | 'Other | 0 | 0 | 0 | 0 | 0 | 81 | 0 | 0 | 0 | 0 |
| 'Clostridiales | 'Ruminococcaceae | 'Oscillospira | '_ | 0 | 0 | 9 | 0 | 34 | 0 | 13 | 12 | 12 | 0 |
| 'Clostridiales | '_ | '_ | '_ | 0 | 0 | 0 | 0 | 0 | 0 | 0 | 0 | 80 | 0 |
| 'Clostridiales | 'Ruminococcaceae | 'Other | 'Other | 8 | 4 | 12 | 0 | 15 | 0 | 25 | 0 | 14 | 0 |
| 'Clostridiales | '_ | '_ | '_ | 0 | 0 | 19 | 0 | 58 | 0 | 0 | 0 | 0 | 0 |
| 'Clostridiales | 'Lachnospiraceae | 'Other | 'Other | 15 | 0 | 0 | 18 | 0 | 0 | 0 | 43 | 0 | 0 |
| 'Clostridiales | 'Other | 'Other | 'Other | 0 | 0 | 0 | 0 | 76 | 0 | 0 | 0 | 0 | 0 |
| 'Lactobacillales | 'Enterococcaceae | 'Vagococcus | '_ | 29 | 0 | 0 | 0 | 0 | 0 | 20 | 0 | 0 | 26 |
| 'Clostridiales | 'Ruminococcaceae | 'Ruminococcus | '_ | 14 | 0 | 9 | 0 | 8 | 0 | 6 | 11 | 23 | 0 |
| 'Erysipelotrichales | 'Erysipelotrichaceae | 'Coprobacillus | '_ | 11 | 0 | 23 | 0 | 0 | 0 | 26 | 0 | 11 | 0 |
| 'Clostridiales | 'Other | 'Other | 'Other | 0 | 0 | 22 | 0 | 49 | 0 | 0 | 0 | 0 | 0 |
| 'Clostridiales | 'Ruminococcaceae | 'Oscillospira | '_ | 0 | 0 | 9 | 0 | 21 | 0 | 14 | 7 | 11 | 9 |
| 'Clostridiales | 'Ruminococcaceae | 'Oscillospira | '_ | 0 | 0 | 38 | 0 | 32 | 0 | 0 | 0 | 0 | 0 |
| 'Clostridiales | '_ | '_ | '_ | 0 | 0 | 14 | 0 | 43 | 0 | 13 | 0 | 0 | 0 |
| 'Clostridiales | 'Lachnospiraceae | '_ | '_ | 0 | 15 | 6 | 14 | 8 | 8 | 8 | 11 | 0 | 0 |
| 'Clostridiales | 'Ruminococcaceae | 'Oscillospira | '_ | 0 | 0 | 0 | 0 | 0 | 0 | 0 | 38 | 0 | 31 |
| 'Clostridiales | 'Lachnospiraceae | 'Coprococcus | '_ | 0 | 0 | 0 | 0 | 52 | 0 | 15 | 0 | 0 | 0 |
| 'Clostridiales | 'Ruminococcaceae | 'Other | 'Other | 0 | 10 | 0 | 9 | 4 | 0 | 7 | 23 | 8 | 5 |
| 'Clostridiales | '_ | '_ | '_ | 0 | 0 | 0 | 0 | 0 | 0 | 26 | 40 | 0 | 0 |
| 'Erysipelotrichales | 'Erysipelotrichaceae | '_ | '_ | 0 | 0 | 9 | 14 | 14 | 14 | 6 | 0 | 6 | 2 |
| 'Bacteroidales | 'Rikenellaceae | '_ | '_ | 0 | 8 | 7 | 4 | 0 | 13 | 7 | 13 | 0 | 11 |
| 'Clostridiales | '_ | '_ | '_ | 0 | 0 | 0 | 0 | 63 | 0 | 0 | 0 | 0 | 0 |
| 'Clostridiales | '_ | '_ | '_ | 0 | 0 | 0 | 0 | 0 | 0 | 21 | 0 | 0 | 40 |
| 'Clostridiales | '_ | '_ | '_ | 15 | 0 | 11 | 0 | 26 | 0 | 0 | 0 | 8 | 0 |
| 'Clostridiales | 'Lachnospiraceae | '_ | '_ | 0 | 0 | 0 | 0 | 58 | 0 | 0 | 0 | 0 | 0 |
| 'Clostridiales | '_ | '_ | '_ | 0 | 0 | 10 | 0 | 32 | 0 | 0 | 0 | 15 | 0 |
| 'Clostridiales | 'Lachnospiraceae | 'Clostridium | 'Other | 0 | 0 | 0 | 0 | 0 | 29 | 0 | 0 | 0 | 28 |
| 'Erysipelotrichales | 'Erysipelotrichaceae | '_ | '_ | 7 | 0 | 18 | 9 | 10 | 0 | 0 | 0 | 9 | 0 |
| 'Lactobacillales | 'Streptococcaceae | 'Streptococcus | 'minor | 0 | 0 | 9 | 0 | 19 | 0 | 16 | 0 | 9 | 0 |
| 'Clostridiales | 'Ruminococcaceae | 'Oscillospira | '_ | 0 | 0 | 0 | 0 | 53 | 0 | 0 | 0 | 0 | 0 |
| 'Clostridiales | '_ | '_ | '_ | 0 | 0 | 0 | 8 | 18 | 0 | 0 | 0 | 27 | 0 |
| 'Clostridiales | 'Lachnospiraceae | 'Other | 'Other | 13 | 0 | 7 | 0 | 13 | 0 | 11 | 0 | 7 | 0 |
| 'Clostridiales | 'Other | 'Other | 'Other | 0 | 0 | 0 | 0 | 15 | 0 | 0 | 0 | 36 | 0 |
| 'RF39 | '_ | '_ | '_ | 0 | 0 | 0 | 0 | 50 | 0 | 0 | 0 | 0 | 0 |
| 'Clostridiales | '_ | '_ | '_ | 0 | 0 | 0 | 0 | 49 | 0 | 0 | 0 | 0 | 0 |
| 'Clostridiales | '_ | '_ | '_ | 14 | 0 | 0 | 0 | 0 | 0 | 0 | 33 | 0 | 0 |
| 'Clostridiales | '_ | '_ | '_ | 0 | 0 | 13 | 0 | 7 | 0 | 0 | 18 | 8 | 0 |
| 'Clostridiales | 'Lachnospiraceae | '_ | '_ | 0 | 11 | 0 | 0 | 0 | 7 | 7 | 0 | 7 | 12 |
| 'Lactobacillales | 'Streptococcaceae | 'Streptococcus | '_ | 0 | 0 | 0 | 0 | 0 | 44 | 0 | 0 | 0 | 0 |
| 'Clostridiales | '_ | '_ | '_ | 0 | 0 | 0 | 0 | 43 | 0 | 0 | 0 | 0 | 0 |
| 'Bacteroidales | 'S24-7 | '_ | '_ | 0 | 0 | 0 | 0 | 0 | 0 | 0 | 43 | 0 | 0 |
| 'Clostridiales | '_ | '_ | '_ | 0 | 0 | 0 | 0 | 0 | 0 | 0 | 0 | 42 | 0 |
| 'Clostridiales | 'Other | 'Other | 'Other | 0 | 0 | 0 | 0 | 0 | 0 | 0 | 0 | 0 | 42 |
| 'Clostridiales | '[Mogibacteriaceae] | '_ | '_ | 7 | 0 | 4 | 12 | 0 | 0 | 6 | 12 | 0 | 0 |
| 'Enterobacteriales | 'Enterobacteriaceae | 'Enterobacter | 'Other | 13 | 0 | 0 | 0 | 0 | 0 | 0 | 27 | 0 | 0 |
| 'Clostridiales | 'Eubacteriaceae | 'Anaerofustis | '_ | 0 | 2 | 15 | 0 | 13 | 0 | 0 | 4 | 5 | 0 |
| 'Erysipelotrichales | 'Erysipelotrichaceae | '_ | '_ | 0 | 0 | 0 | 0 | 15 | 0 | 5 | 0 | 0 | 19 |
| 'Lactobacillales | 'Aerococcaceae | 'Aerococcus | '_ | 0 | 0 | 0 | 0 | 0 | 0 | 0 | 0 | 0 | 39 |
| 'Clostridiales | 'Ruminococcaceae | 'Oscillospira | '_ | 0 | 0 | 21 | 0 | 17 | 0 | 0 | 0 | 0 | 0 |
| 'Clostridiales | 'Ruminococcaceae | 'Oscillospira | '_ | 0 | 0 | 0 | 10 | 0 | 0 | 0 | 16 | 0 | 12 |
| 'Clostridiales | 'Other | 'Other | 'Other | 0 | 0 | 16 | 0 | 9 | 0 | 7 | 0 | 5 | 0 |
| 'Clostridiales | '_ | '_ | '_ | 0 | 9 | 0 | 10 | 0 | 5 | 0 | 0 | 5 | 8 |
| 'Clostridiales | 'Lachnospiraceae | 'Coprococcus | '_ | 0 | 0 | 17 | 0 | 0 | 0 | 0 | 18 | 0 | 0 |
| 'Clostridiales | 'Other | 'Other | 'Other | 0 | 0 | 0 | 0 | 0 | 0 | 0 | 0 | 35 | 0 |
| 'Clostridiales | '_ | '_ | '_ | 0 | 0 | 33 | 0 | 0 | 0 | 0 | 0 | 0 | 0 |
| 'Clostridiales | 'Lachnospiraceae | 'Other | 'Other | 0 | 0 | 12 | 0 | 0 | 0 | 9 | 0 | 11 | 0 |
| 'Clostridiales | 'Ruminococcaceae | 'Oscillospira | '_ | 0 | 0 | 8 | 14 | 0 | 0 | 0 | 10 | 0 | 0 |
| 'Bacteroidales | 'S24-7 | '_ | '_ | 0 | 32 | 0 | 0 | 0 | 0 | 0 | 0 | 0 | 0 |
| 'Clostridiales | 'Ruminococcaceae | 'Ruminococcus | '_ | 0 | 0 | 15 | 0 | 16 | 0 | 0 | 0 | 0 | 0 |
| 'Clostridiales | 'Ruminococcaceae | 'Oscillospira | '_ | 0 | 0 | 0 | 0 | 0 | 0 | 0 | 0 | 31 | 0 |
| 'Clostridiales | '_ | '_ | '_ | 0 | 0 | 30 | 0 | 0 | 0 | 0 | 0 | 0 | 0 |
| 'Clostridiales | '_ | '_ | '_ | 0 | 0 | 14 | 0 | 0 | 0 | 16 | 0 | 0 | 0 |
| 'Clostridiales | '_ | '_ | '_ | 0 | 0 | 0 | 0 | 30 | 0 | 0 | 0 | 0 | 0 |
| 'Clostridiales | '_ | '_ | '_ | 0 | 0 | 0 | 8 | 6 | 0 | 0 | 0 | 16 | 0 |
| 'Clostridiales | 'Ruminococcaceae | 'Ruminococcus | '_ | 0 | 12 | 0 | 8 | 0 | 0 | 0 | 10 | 0 | 0 |
| 'Clostridiales | 'Lachnospiraceae | '_ | '_ | 6 | 0 | 7 | 0 | 6 | 0 | 0 | 10 | 0 | 0 |
| 'Clostridiales | 'Ruminococcaceae | 'Oscillospira | '_ | 0 | 0 | 14 | 0 | 0 | 0 | 15 | 0 | 0 | 0 |
| 'Clostridiales | '_ | '_ | '_ | 0 | 0 | 0 | 0 | 15 | 0 | 0 | 0 | 14 | 0 |
| 'Clostridiales | '_ | '_ | '_ | 0 | 0 | 0 | 0 | 0 | 0 | 9 | 0 | 20 | 0 |
| 'Actinomycetales | 'Corynebacteriaceae | 'Corynebacterium | 'stationis | 26 | 0 | 0 | 0 | 0 | 0 | 0 | 0 | 0 | 0 |
| 'Clostridiales | 'Ruminococcaceae | 'Other | 'Other | 0 | 0 | 4 | 0 | 3 | 0 | 14 | 0 | 5 | 0 |
| 'Bacteroidales | 'S24-7 | '_ | '_ | 0 | 0 | 0 | 0 | 0 | 0 | 0 | 13 | 0 | 13 |
| 'Clostridiales | 'Ruminococcaceae | 'Oscillospira | '_ | 0 | 0 | 0 | 0 | 0 | 0 | 0 | 0 | 0 | 26 |
| 'Clostridiales | 'Ruminococcaceae | 'Oscillospira | '_ | 0 | 0 | 0 | 0 | 0 | 0 | 0 | 25 | 0 | 0 |
| 'Clostridiales | 'Ruminococcaceae | 'Ruminococcus | '_ | 13 | 0 | 11 | 0 | 0 | 0 | 0 | 0 | 0 | 0 |
| 'Clostridiales | '[Mogibacteriaceae] | '_ | '_ | 11 | 0 | 0 | 4 | 0 | 0 | 0 | 9 | 0 | 0 |
| 'Erysipelotrichales | 'Erysipelotrichaceae | 'Coprobacillus | '_ | 0 | 0 | 0 | 0 | 14 | 0 | 10 | 0 | 0 | 0 |
| 'Erysipelotrichales | 'Erysipelotrichaceae | 'Clostridium | 'cocleatum | 0 | 2 | 0 | 0 | 0 | 0 | 0 | 20 | 0 | 1 |
| 'Bacteroidales | 'S24-7 | '_ | '_ | 0 | 0 | 0 | 0 | 0 | 0 | 0 | 23 | 0 | 0 |
| 'Coriobacteriales | 'Coriobacteriaceae | '_ | '_ | 0 | 0 | 7 | 0 | 10 | 5 | 0 | 0 | 0 | 0 |
| 'Coriobacteriales | 'Coriobacteriaceae | 'Adlercreutzia | '_ | 4 | 0 | 0 | 0 | 8 | 9 | 0 | 0 | 0 | 0 |
| 'Lactobacillales | 'Leuconostocaceae | 'Weissella | 'paramesenteroides | 2 | 0 | 0 | 3 | 0 | 0 | 4 | 0 | 0 | 12 |
| 'Lactobacillales | 'Enterococcaceae | 'Vagococcus | '_ | 0 | 0 | 0 | 0 | 21 | 0 | 0 | 0 | 0 | 0 |
| 'Clostridiales | 'Ruminococcaceae | 'Butyricicoccus | 'pullicaecorum | 0 | 0 | 0 | 0 | 7 | 0 | 0 | 8 | 6 | 0 |
| 'Clostridiales | '[Mogibacteriaceae] | '_ | '_ | 0 | 0 | 0 | 12 | 0 | 0 | 0 | 3 | 0 | 6 |
| 'RF39 | '_ | '_ | '_ | 0 | 0 | 0 | 0 | 12 | 0 | 0 | 0 | 8 | 0 |
| 'Clostridiales | '_ | '_ | '_ | 0 | 0 | 0 | 0 | 0 | 0 | 0 | 0 | 20 | 0 |
| 'Clostridiales | '_ | '_ | '_ | 0 | 0 | 0 | 0 | 0 | 0 | 0 | 0 | 20 | 0 |
| 'Bacteroidales | 'S24-7 | '_ | '_ | 0 | 0 | 0 | 9 | 0 | 0 | 0 | 0 | 0 | 11 |
| 'Clostridiales | 'Ruminococcaceae | 'Oscillospira | '_ | 0 | 0 | 9 | 0 | 0 | 0 | 0 | 10 | 0 | 0 |
| 'Bacillales | 'Planococcaceae | 'Sporosarcina | '_ | 0 | 8 | 0 | 0 | 3 | 3 | 4 | 0 | 0 | 0 |
| 'Clostridiales | 'Lachnospiraceae | 'Blautia | 'producta | 0 | 0 | 7 | 6 | 0 | 0 | 4 | 0 | 0 | 0 |
| 'Erysipelotrichales | 'Erysipelotrichaceae | 'Other | 'Other | 0 | 0 | 0 | 0 | 0 | 10 | 0 | 0 | 7 | 0 |
| 'Clostridiales | '_ | '_ | '_ | 0 | 0 | 0 | 0 | 0 | 0 | 16 | 0 | 0 | 0 |
| 'Clostridiales | 'Ruminococcaceae | 'Ruminococcus | '_ | 0 | 0 | 15 | 0 | 0 | 0 | 0 | 0 | 0 | 0 |
| 'Clostridiales | '_ | '_ | '_ | 0 | 0 | 0 | 0 | 15 | 0 | 0 | 0 | 0 | 0 |
| 'Clostridiales | '_ | '_ | '_ | 0 | 0 | 0 | 0 | 0 | 0 | 15 | 0 | 0 | 0 |
| 'Clostridiales | '_ | '_ | '_ | 0 | 15 | 0 | 0 | 0 | 0 | 0 | 0 | 0 | 0 |
| 'Coriobacteriales | 'Coriobacteriaceae | 'Adlercreutzia | '_ | 10 | 4 | 0 | 0 | 0 | 0 | 0 | 0 | 0 | 0 |
| 'Clostridiales | 'Other | 'Other | 'Other | 0 | 0 | 0 | 0 | 0 | 0 | 14 | 0 | 0 | 0 |
| 'Clostridiales | 'Ruminococcaceae | 'Other | 'Other | 0 | 5 | 0 | 9 | 0 | 0 | 0 | 0 | 0 | 0 |
| 'Erysipelotrichales | 'Erysipelotrichaceae | '_ | '_ | 0 | 0 | 0 | 14 | 0 | 0 | 0 | 0 | 0 | 0 |
| 'Deferribacterales | 'Deferribacteraceae | 'Mucispirillum | 'schaedleri | 0 | 0 | 0 | 0 | 0 | 14 | 0 | 0 | 0 | 0 |
| 'Rhizobiales | 'Methylobacteriaceae | 'Methylobacterium | 'Other | 0 | 0 | 0 | 0 | 2 | 11 | 0 | 0 | 0 | 0 |
| 'Methanobacteriales | 'Methanobacteriaceae | 'Methanobrevibacter | '_ | 0 | 0 | 0 | 0 | 0 | 13 | 0 | 0 | 0 | 0 |
| 'RF39 | '_ | '_ | '_ | 0 | 2 | 6 | 0 | 4 | 0 | 0 | 0 | 0 | 0 |
| 'Clostridiales | 'Lachnospiraceae | 'Other | 'Other | 0 | 0 | 0 | 0 | 0 | 0 | 0 | 0 | 12 | 0 |
| 'Clostridiales | 'Lachnospiraceae | 'Dorea | '_ | 11 | 0 | 0 | 0 | 0 | 0 | 0 | 0 | 0 | 0 |
| 'Clostridiales | 'Lachnospiraceae | 'Dorea | '_ | 0 | 0 | 4 | 0 | 0 | 0 | 0 | 0 | 7 | 0 |
| 'Clostridiales | 'Lachnospiraceae | '_ | '_ | 0 | 0 | 0 | 0 | 0 | 0 | 0 | 0 | 11 | 0 |
| 'Erysipelotrichales | 'Erysipelotrichaceae | '_ | '_ | 0 | 11 | 0 | 0 | 0 | 0 | 0 | 0 | 0 | 0 |
| 'Clostridiales | '_ | '_ | '_ | 0 | 11 | 0 | 0 | 0 | 0 | 0 | 0 | 0 | 0 |
| 'Clostridiales | 'Other | 'Other | 'Other | 0 | 0 | 0 | 0 | 0 | 0 | 0 | 11 | 0 | 0 |
| 'Clostridiales | 'Ruminococcaceae | 'Ruminococcus | '_ | 0 | 0 | 0 | 0 | 0 | 0 | 0 | 0 | 0 | 11 |
| 'Clostridiales | 'Lachnospiraceae | 'Other | 'Other | 0 | 0 | 10 | 0 | 0 | 0 | 0 | 0 | 0 | 0 |
| 'Erysipelotrichales | 'Erysipelotrichaceae | 'Clostridium | 'cocleatum | 0 | 0 | 0 | 0 | 10 | 0 | 0 | 0 | 0 | 0 |
| 'Clostridiales | 'Clostridiaceae | '_ | '_ | 0 | 0 | 0 | 0 | 0 | 0 | 0 | 0 | 10 | 0 |
| 'Bacteroidales | 'S24-7 | '_ | '_ | 0 | 0 | 0 | 6 | 0 | 0 | 0 | 4 | 0 | 0 |
| 'Clostridiales | 'Ruminococcaceae | 'Oscillospira | '_ | 0 | 0 | 0 | 0 | 0 | 0 | 0 | 0 | 0 | 10 |
| 'Clostridiales | '_ | '_ | '_ | 9 | 0 | 0 | 0 | 0 | 0 | 0 | 0 | 0 | 0 |
| 'Clostridiales | '_ | '_ | '_ | 0 | 0 | 0 | 0 | 9 | 0 | 0 | 0 | 0 | 0 |
| 'Erysipelotrichales | 'Erysipelotrichaceae | '_ | '_ | 0 | 0 | 0 | 0 | 0 | 0 | 0 | 0 | 0 | 9 |
| 'Lactobacillales | 'Streptococcaceae | '_ | '_ | 0 | 0 | 0 | 0 | 8 | 0 | 0 | 0 | 0 | 0 |
| 'Erysipelotrichales | 'Erysipelotrichaceae | '_ | '_ | 0 | 0 | 0 | 0 | 0 | 0 | 8 | 0 | 0 | 0 |
| 'Turicibacterales | 'Turicibacteraceae | 'Turicibacter | '_ | 7 | 0 | 0 | 0 | 0 | 0 | 0 | 0 | 0 | 0 |
| 'Bacteroidales | 'S24-7 | '_ | '_ | 0 | 0 | 7 | 0 | 0 | 0 | 0 | 0 | 0 | 0 |
| 'Clostridiales | 'Ruminococcaceae | 'Oscillospira | '_ | 0 | 0 | 7 | 0 | 0 | 0 | 0 | 0 | 0 | 0 |
| 'Clostridiales | 'Lachnospiraceae | '_ | '_ | 0 | 0 | 7 | 0 | 0 | 0 | 0 | 0 | 0 | 0 |
| 'Clostridiales | '_ | '_ | '_ | 0 | 3 | 0 | 0 | 0 | 0 | 0 | 0 | 0 | 4 |
| 'Erysipelotrichales | 'Erysipelotrichaceae | '_ | '_ | 0 | 0 | 0 | 0 | 0 | 0 | 0 | 7 | 0 | 0 |
| 'Clostridiales | 'Ruminococcaceae | 'Ruminococcus | '_ | 0 | 0 | 0 | 0 | 0 | 0 | 0 | 0 | 0 | 7 |
| 'Pseudomonadales | 'Moraxellaceae | 'Psychrobacter | 'Other | 3 | 0 | 0 | 0 | 0 | 0 | 3 | 0 | 0 | 0 |
| 'Bacillales | 'Paenibacillaceae | 'Other | 'Other | 0 | 0 | 6 | 0 | 0 | 0 | 0 | 0 | 0 | 0 |
| 'Clostridiales | 'Lachnospiraceae | 'Coprococcus | '_ | 0 | 0 | 6 | 0 | 0 | 0 | 0 | 0 | 0 | 0 |
| 'Bacillales | 'Bacillaceae | 'Bacillus | '_ | 0 | 0 | 0 | 0 | 6 | 0 | 0 | 0 | 0 | 0 |
| 'Lactobacillales | 'Enterococcaceae | 'Vagococcus | '_ | 0 | 0 | 0 | 0 | 0 | 0 | 6 | 0 | 0 | 0 |
| 'RF39 | '_ | '_ | '_ | 0 | 0 | 0 | 0 | 0 | 6 | 0 | 0 | 0 | 0 |
| 'Clostridiales | 'Lachnospiraceae | '_ | '_ | 0 | 0 | 0 | 0 | 0 | 0 | 0 | 6 | 0 | 0 |
| 'Clostridiales | '_ | '_ | '_ | 0 | 0 | 0 | 0 | 0 | 0 | 0 | 0 | 0 | 6 |
| 'RF39 | '_ | '_ | '_ | 5 | 0 | 0 | 0 | 0 | 0 | 0 | 0 | 0 | 0 |
| 'Clostridiales | '_ | '_ | '_ | 0 | 0 | 0 | 0 | 0 | 0 | 0 | 0 | 5 | 0 |
| 'Clostridiales | 'Ruminococcaceae | 'Oscillospira | '_ | 0 | 0 | 4 | 0 | 0 | 0 | 0 | 0 | 0 | 0 |
| 'RF39 | '_ | '_ | '_ | 0 | 0 | 0 | 0 | 4 | 0 | 0 | 0 | 0 | 0 |
| 'Erysipelotrichales | 'Erysipelotrichaceae | '_ | '_ | 0 | 0 | 0 | 0 | 0 | 0 | 0 | 0 | 4 | 0 |
| 'Clostridiales | 'Other | 'Other | 'Other | 0 | 0 | 0 | 0 | 0 | 0 | 0 | 0 | 0 | 4 |
| 'Clostridiales | 'Clostridiaceae | 'Clostridium | 'Other | 3 | 0 | 0 | 0 | 0 | 0 | 0 | 0 | 0 | 0 |
| 'Clostridiales | '_ | '_ | '_ | 0 | 0 | 0 | 0 | 3 | 0 | 0 | 0 | 0 | 0 |
| 'Clostridiales | 'Ruminococcaceae | 'Butyricicoccus | 'pullicaecorum | 0 | 0 | 0 | 0 | 0 | 0 | 3 | 0 | 0 | 0 |
| 'Clostridiales | 'Ruminococcaceae | '_ | '_ | 0 | 0 | 0 | 0 | 0 | 0 | 0 | 0 | 3 | 0 |
| 'Deferribacterales | 'Deferribacteraceae | 'Mucispirillum | 'schaedleri | 0 | 0 | 0 | 3 | 0 | 0 | 0 | 0 | 0 | 0 |
| 'Turicibacterales | 'Turicibacteraceae | 'Turicibacter | '_ | 0 | 0 | 0 | 0 | 0 | 0 | 0 | 3 | 0 | 0 |
| 'Clostridiales | '_ | '_ | '_ | 0 | 0 | 0 | 0 | 0 | 0 | 0 | 0 | 0 | 3 |
| 'Verrucomicrobiales | 'Verrucomicrobiaceae | 'Akkermansia | 'muciniphila | 2 | 0 | 0 | 0 | 0 | 0 | 0 | 0 | 0 | 0 |
| 'Clostridiales | '[Mogibacteriaceae] | '_ | '_ | 0 | 0 | 2 | 0 | 0 | 0 | 0 | 0 | 0 | 0 |
| 'Burkholderiales | 'Alcaligenaceae | 'Sutterella | '_ | 0 | 0 | 0 | 0 | 0 | 0 | 2 | 0 | 0 | 0 |
| 'Clostridiales | 'Lachnospiraceae | 'Dorea | 'longicatena | 0 | 0 | 0 | 0 | 0 | 0 | 0 | 0 | 2 | 0 |
| 'Clostridiales | 'Ruminococcaceae | '_ | '_ | 0 | 0 | 0 | 2 | 0 | 0 | 0 | 0 | 0 | 0 |
| 'Clostridiales | 'Lachnospiraceae | '_ | '_ | 0 | 0 | 0 | 0 | 0 | 2 | 0 | 0 | 0 | 0 |
| 'Clostridiales | '_ | '_ | '_ | 0 | 0 | 0 | 0 | 0 | 0 | 0 | 0 | 0 | 2 |

**Supplementary Table 2:** Peak height of all the detected metabolite compounds after normalization

| **S.No** | **Name of the Compound** | **Gr.1 - Control** | | | **Gr.2 -AFO-202** | | | **Gr. 3- N-163** | | | **Gr.4 - AFO-202+N-163** | | | **Telmisartan** | | |
| --- | --- | --- | --- | --- | --- | --- | --- | --- | --- | --- | --- | --- | --- | --- | --- | --- |
|  |  | **Pre** | **Post** | **Difference** | **Pre** | **Post** | **Difference** | **Pre** | **Post** | **Difference** | **Pre** | **Post** | **Difference** | **Pre** | **Post** | **Difference** |
| 1 | 2-Aminobutyric acid-2TMS | 16544.74 | 40209.97 | 23665.23 | 23704.19 | 27627.27 | 3923.08 | 11065.45 | 24994.24 | 13928.79 | 24652.04 | 39540.28 | 14888.24 | 25746.06 | 43700.57 | 17954.52 |
| 2 | 2-Hydroxyisobutyric acid-2TMS | 36888.72 | 145407.17 | 108518.45 | 38444.81 | 146808.33 | 108363.52 | 70728.84 | 162704.66 | 91975.82 | 38044.04 | 179597.18 | 141553.15 | 36852.36 | 170732.73 | 133880.36 |
| 3 | 3-Aminoglutaric acid-3TMS | 2064.63 | 11599.10 | 9534.47 | 2674.96 | 8297.81 | 5622.84 | 2061.59 | 4755.57 | 2693.99 | 4899.49 | 5638.95 | 739.46 | 3808.51 | 12436.04 | 8627.53 |
| 4 | 3-Hydroxybutyric acid-2TMS | 78693.42 | 32663.28 | **-46030.14** | 49425.90 | 15869.18 | **-33556.73** | 59399.24 | 16902.03 | **-42497.20** | 23465.34 | 16080.23 | **-7385.11** | 15499.52 | 10891.34 | **-4608.17** |
| 5 | 3-Methyl-2-oxovaleric acid-meto-TMS | 2494.33 | 15765.26 | 13270.93 | 5771.05 | 12154.53 | 6383.48 | 3664.87 | 10069.54 | 6404.67 | 6043.01 | 13013.54 | 6970.53 | 7152.49 | 12393.37 | 5240.87 |
| 6 | 4-Hydroxyphenylacetic acid-2TMS | 71212.24 | 90389.17 | 19176.93 | 38406.22 | 61406.95 | 23000.73 | 81498.41 | 108347.44 | 26849.04 | 29763.24 | 65944.94 | 36181.70 | 36067.25 | 39882.54 | 3815.29 |
| 7 | 5-Aminovaleric acid-3TMS | 359981.32 | 177455.86 | **-182525.46** | 28606.18 | 220398.95 | 191792.77 | 184342.02 | 312636.97 | 128294.95 | 435424.34 | 319991.71 | **-115432.64** | 28517.91 | 292458.93 | 263941.02 |
| 8 | 5-Oxoproline-2TMS | 86202.33 | 183221.07 | 97018.75 | 101052.25 | 142117.70 | 41065.45 | 76536.42 | 135748.92 | 59212.51 | 74316.31 | 137827.60 | 63511.29 | 113169.35 | 198991.94 | 85822.59 |
| 9 | Acetylglycine-TMS | 62276.84 | 78877.14 | 16600.30 | 62217.09 | 75225.03 | 13007.93 | 54550.93 | 64278.24 | 9727.30 | 47481.52 | 68504.55 | 21023.03 | 70946.14 | 99435.23 | 28489.09 |
| 10 | Alanine-2TMS | 462486.71 | 932281.83 | 469795.13 | 575514.07 | 709299.17 | 133785.10 | 381431.60 | 576714.21 | 195282.60 | 451475.22 | 778523.96 | 327048.74 | 544032.57 | 1268525.27 | 724492.70 |
| 11 | Asparagine-3TMS | 50264.14 | 47387.02 | **-2877.12** | 52065.54 | 39742.05 | **-12323.49** | 45670.67 | 23805.15 | **-21865.52** | 40462.65 | 35329.03 | **-5133.62** | 57492.79 | 68357.27 | 10864.48 |
| 12 | Aspartic acid-3TMS | 122316.67 | 309390.39 | 187073.73 | 168305.85 | 250489.61 | 82183.75 | 103731.51 | 227874.35 | 124142.84 | 133017.58 | 208828.93 | 75811.35 | 183416.60 | 433233.75 | 249817.15 |
| 13 | Fructose-meto-5TMS | ######### | 999935.86 | **-726073.07** | 323320.36 | 936708.26 | 613387.90 | 2412757.75 | 1126616.20 | **-1286141.55** | 1689190.48 | 2733495.73 | 1044305.25 | 325202.45 | 1110939.22 | 785736.77 |
| 14 | Fucose-meto-4TMS | 36824.30 | 44185.98 | 7361.68 | 57488.33 | 33927.51 | **-23560.83** | 58516.62 | 21689.03 | **-36827.59** | 58635.69 | 53958.65 | **-4677.04** | 89055.59 | 24027.13 | **-65028.47** |
| 15 | Galactose-meto-5TMS | 92513.44 | 457335.67 | 364822.23 | 177745.54 | 661315.76 | 483570.22 | 191900.52 | 559295.50 | 367394.98 | 139909.99 | 1341594.81 | 1201684.82 | 359608.12 | 188781.00 | **-170827.12** |
| 16 | Glucose-meto-5TMS | 299911.54 | 1740399.37 | 1440487.83 | 842456.89 | 996893.55 | 154436.66 | 448031.24 | 1023564.45 | 575533.20 | 273170.40 | 1501256.09 | 1228085.70 | 400546.16 | 855701.34 | 455155.19 |
| 17 | Glutamic acid-3TMS | 197256.75 | 658386.79 | 461130.05 | 264245.78 | 515323.80 | 251078.03 | 170970.32 | 416395.95 | 245425.63 | 269851.71 | 394663.22 | 124811.50 | 361041.65 | 743751.06 | 382709.41 |
| 18 | Glutamine-3TMS | 60900.94 | 73851.25 | 12950.31 | 45289.87 | 46071.10 | 781.23 | 40503.29 | 47109.70 | 6606.41 | 27222.10 | 52375.70 | 25153.60 | 46899.14 | 77644.26 | 30745.13 |
| 19 | Glycerol-3TMS | 100045.16 | 512115.32 | 412070.17 | 193048.44 | 332266.25 | 139217.82 | 351596.87 | 290395.66 | **-61201.21** | 124107.19 | 374822.76 | 250715.57 | 283873.86 | 324467.91 | 40594.06 |
| 20 | Glycine-3TMS | 175941.04 | 190667.24 | 14726.20 | 182268.66 | 182802.22 | 533.56 | 118303.95 | 134185.05 | 15881.10 | 148645.59 | 111069.96 | **-37575.63** | 179124.85 | 288456.14 | 109331.29 |
| 21 | Hypoxanthine-2TMS | 18209.77 | 73610.39 | 55400.63 | 25970.42 | 54650.62 | 28680.21 | 23749.72 | 48430.05 | 24680.32 | 22627.37 | 60014.19 | 37386.83 | 33144.25 | 56824.97 | 23680.72 |
| 22 | Inosine-4TMS | 20341.44 | 29963.22 | 9621.79 | 27134.09 | 52940.46 | 25806.38 | 27591.81 | 56079.56 | 28487.75 | 33777.00 | 42475.35 | 8698.35 | 24548.88 | 37226.82 | 12677.95 |
| 23 | Isoleucine-2TMS | 240751.69 | 255679.87 | 14928.19 | 213617.13 | 185078.81 | **-28538.33** | 142467.71 | 156224.53 | 13756.81 | 140230.24 | 158650.59 | 18420.35 | 175687.57 | 381173.16 | 205485.59 |
| 24 | Lactic acid-2TMS | 126966.77 | 174366.10 | 47399.33 | 161320.99 | 112769.78 | **-48551.21** | 181299.93 | 76773.16 | **-104526.77** | 52406.61 | 197790.91 | 145384.31 | 71969.55 | 36363.66 | **-35605.88** |
| 25 | Leucine-2TMS | 405177.22 | 484969.86 | 79792.64 | 381285.89 | 366743.37 | **-14542.52** | 297859.97 | 296018.19 | **-1841.78** | 273676.66 | 287240.08 | 13563.42 | 364222.50 | 631954.96 | 267732.46 |
| 26 | Lysine-4TMS | 453167.29 | 1042910.27 | 589742.97 | 578069.23 | 771280.17 | 193210.94 | 495585.73 | 790010.41 | 294424.68 | 505753.28 | 831999.15 | 326245.87 | 592034.86 | 952876.14 | 360841.28 |
| 27 | Malic acid-3TMS | 15431.92 | 50833.82 | 35401.90 | 18557.13 | 136072.04 | 117514.91 | 23517.14 | 116959.69 | 93442.55 | 25340.81 | 67836.64 | 42495.84 | 54899.38 | 38544.73 | **-16354.66** |
| 28 | Mannose-meto-5TMS | 46604.74 | 101647.05 | 55042.30 | 58090.57 | 125118.62 | 67028.06 | 70844.17 | 147959.66 | 77115.49 | 45441.01 | 167518.11 | 122077.10 | 79951.11 | 65364.10 | **-14587.00** |
| 29 | Methionine-2TMS | 72809.85 | 128516.84 | 55706.99 | 84612.94 | 90620.86 | 6007.93 | 66116.46 | 88928.83 | 22812.37 | 68604.44 | 92418.75 | 23814.31 | 103825.98 | 168797.44 | 64971.46 |
| 30 | N-Acetylmannosamine-meto-4TMS | 51349.02 | 194543.70 | 143194.68 | 95958.86 | 210636.96 | 114678.10 | 102946.69 | 130222.12 | 27275.43 | 102728.94 | 270379.17 | 167650.23 | 207515.87 | 90678.54 | **-116837.33** |
| 31 | Nicotinic acid-TMS | 8270.56 | 31967.10 | 23696.54 | 12163.20 | 23936.48 | 11773.28 | 11141.64 | 22880.62 | 11738.98 | 12707.43 | 23489.75 | 10782.32 | 18359.32 | 23826.50 | 5467.18 |
| 32 | Norvaline-TMS | 94923.32 | 100191.34 | 5268.03 | 82912.31 | 73583.61 | **-9328.70** | 58328.64 | 66992.36 | 8663.72 | 54802.10 | 77669.25 | 22867.15 | 72621.50 | 110365.67 | 37744.18 |
| 33 | Ornithine-4TMS | 45698.47 | 101711.56 | 56013.10 | 40654.29 | 52324.86 | 11670.57 | 52234.33 | 56216.15 | 3981.82 | 54989.59 | 101736.85 | 46747.26 | 44496.80 | 62226.84 | 17730.04 |
| 34 | Phenylalanine-2TMS | 173673.02 | 290028.18 | 116355.17 | 186236.03 | 228284.46 | 42048.43 | 161361.25 | 202463.07 | 41101.82 | 145354.48 | 179722.22 | 34367.74 | 208441.78 | 342996.16 | 134554.38 |
| 35 | Phosphoric acid-3TMS | 13703.35 | 692470.05 | 678766.69 | 55467.68 | 765279.47 | 709811.79 | 39524.43 | 1030958.21 | 991433.78 | 51518.47 | 879380.28 | 827861.80 | 107073.44 | 122301.92 | 15228.48 |
| 36 | Proline-2TMS | 57195.44 | 92021.21 | 34825.77 | 77037.72 | 103324.97 | 26287.25 | 60257.37 | 83664.37 | 23407.00 | 66855.91 | 53963.25 | **-12892.66** | 98386.02 | 151136.53 | 52750.51 |
| 37 | Putrescine-4TMS | 90940.03 | 4806.17 | **-86133.86** | 30410.15 | 5030.93 | **-25379.22** | 70539.24 | 6011.09 | **-64528.14** | 62824.68 | 9891.53 | **-52933.15** | 21749.83 | 6122.91 | **-15626.91** |
| 38 | Pyruvic acid-meto-TMS | 31557.87 | 108630.52 | 77072.65 | 40255.26 | 105996.98 | 65741.72 | 52793.70 | 107097.52 | 54303.82 | 40456.18 | 121792.59 | 81336.41 | 41703.69 | 116493.60 | 74789.91 |
| 39 | Ribonic acid-5TMS | 90115.51 | 64726.74 | -25388.76 | 85151.16 | 66560.71 | -18590.46 | 90412.13 | 99956.38 | 9544.25 | 93736.17 | 93030.85 | **-705.32** | 95701.95 | 64707.65 | -30994.30 |
| 40 | Ribose-meto-4TMS | 199285.35 | 806414.91 | 607129.56 | 321848.24 | 706173.47 | 384325.23 | 284999.93 | 561917.45 | 276917.52 | 289686.98 | 526890.91 | 237203.94 | 510635.39 | 699650.70 | 189015.32 |
| 41 | Serine-3TMS | 134100.10 | 150072.29 | 15972.19 | 134521.62 | 135009.56 | 487.94 | 94829.48 | 89019.26 | -5810.21 | 87612.04 | 97022.85 | 9410.81 | 124526.34 | 206603.74 | 82077.40 |
| 42 | Spermidine-5TMS | 11571.79 | 14335.92 | 2764.13 | 8257.93 | 7462.86 | **-795.07** | 10699.82 | 7443.06 | **-3256.75** | 10274.90 | 8775.14 | **-1499.76** | 9612.74 | 8122.10 | **-1490.65** |
| 43 | Succinic acid-2TMS | 200720.47 | 75253.68 | **-125466.79** | 32921.32 | 1398291.07 | 1365369.75 | 383811.07 | 1183826.74 | 800015.68 | 44403.33 | 504988.19 | 460584.87 | 263097.91 | 313904.61 | 50806.69 |
| 44 | Taurine-3TMS | 15049.13 | 66033.11 | 50983.98 | 14685.18 | 46705.91 | 32020.73 | 11165.27 | 32313.06 | 21147.79 | 14449.38 | 55683.04 | 41233.65 | 14857.44 | 24726.77 | 9869.33 |
| 45 | Threonine-3TMS | 137361.71 | 175385.78 | 38024.08 | 137475.15 | 135608.93 | **-1866.22** | 91893.96 | 113191.42 | 21297.46 | 87758.08 | 115202.12 | 27444.04 | 122185.79 | 240869.77 | 118683.98 |
| 46 | Thymine-2TMS | 5319.57 | 16561.64 | 11242.07 | 6636.79 | 13272.69 | 6635.90 | 6900.32 | 9466.94 | 2566.62 | 9972.14 | 17972.30 | 8000.16 | 12893.64 | 7133.20 | **-5760.44** |
| 47 | Tryptophan-3TMS | 21044.89 | 20895.79 | **-149.10** | 17365.35 | 18254.73 | 889.38 | 13733.63 | 17318.66 | 3585.03 | 14840.33 | 14286.41 | **-553.92** | 16169.30 | 28678.00 | 12508.70 |
| 48 | Tyrosine-3TMS | 273469.68 | 438287.99 | 164818.32 | 293828.48 | 351487.07 | 57658.59 | 234275.64 | 316786.47 | 82510.82 | 232207.59 | 363478.28 | 131270.69 | 309190.46 | 425825.94 | 116635.49 |
| 49 | Uracil-2TMS | 25078.62 | 92837.69 | 67759.07 | 44546.07 | 84845.85 | 40299.79 | 35546.66 | 66320.55 | 30773.89 | 50653.91 | 65823.53 | 15169.62 | 74843.38 | 71567.16 | **-3276.22** |
| 50 | Valine-2TMS | 353165.15 | 405846.51 | 52681.36 | 290698.34 | 255449.25 | **-35249.08** | 219247.47 | 233471.87 | 14224.40 | 202562.77 | 238231.09 | 35668.32 | 254032.97 | 521841.20 | 267808.23 |
| 51 | Xanthine-3TMS | 53777.98 | 130586.54 | 76808.56 | 79540.73 | 113793.80 | 34253.08 | 83412.44 | 89448.75 | 6036.32 | 71847.21 | 109146.53 | 37299.32 | 101684.60 | 103194.40 | 1509.80 |
| 52 | Xylose-meto-4TMS | 174559.36 | 290187.08 | 115627.72 | 186404.46 | 228374.90 | 41970.44 | 162359.78 | 202528.45 | 40168.67 | 145455.14 | 179768.76 | 34313.61 | 208574.29 | 343176.94 | 134602.65 |

**Supplementary Table 3:** Coefficients of Metabolite compounds with a VIP value of 1 or higher in OPLS-DA in the different groups

| **No.** | **Metabolites** | **Coefficients** | | | | |
| --- | --- | --- | --- | --- | --- | --- |
|  |  | **Control** | **AFO-202** | **N-163** | **AFO-202+N-163** | **Telmisartan** |
| 1 | Glucose-meto-5TMS | -0.000180 | -0.000091 | -0.000151 | -0.000168 | -0.000139 |
| 2 | Fructose-meto-5TMS | 0.000133 | -0.000173 | 0.000215 | -0.000177 | -0.000209 |
| 3 | Phosphoric acid-3TMS | -0.000125 |  |  |  |  |
| 4 | Ribose-meto-4TMS | -0.000116 | -0.000132 | -0.000103 | -0.000065 | -0.000083 |
| 5 | Lysine-4TMS | -0.000116 | -0.000083 | -0.000098 | -0.000069 | -0.000120 |
| 6 | Alanine-2TMS | -0.000109 | -0.000003 | -0.000061 | -0.000094 | -0.000171 |
| 7 | Glutamic acid-3TMS | -0.000100 | -0.000101 | -0.000102 | -0.000063 | -0.000149 |
| 8 | Glycerol-3TMS | -0.000102 | -0.000080 |  | -0.000084 |  |
| 9 | Galactose-meto-5TMS | -0.000092 | -0.000144 | -0.000124 | -0.000190 | 0.000094 |
| 10 | Aspartic acid-3TMS | -0.000063 | -0.000061 | -0.000064 |  | -0.000113 |
| 11 | 5-Aminovaleric acid-3TMS | 0.000069 | -0.000103 | -0.000077 | 0.000073 | -0.000123 |
| 12 | Tyrosine-3TMS | -0.000064 |  | -0.000039 | -0.000057 | -0.000043 |
| 13 | *N*-Acetylmannosamine-meto-4TMS | -0.000055 | -0.000068 |  | -0.000058 |  |
| 14 | Succinic acid-2TMS | 0.000059 | -0.000259 | -0.000172 | -0.000109 |  |
| 15 | Phenylalanine-2TMS | -0.000049 |  |  |  | -0.000087 |
| 16 | Xylose-meto-4TMS | -0.000048 |  |  |  | -0.000087 |
| 17 | 2-Hydroxyisobutyric acid-2TMS | -0.000054 | -0.000072 | -0.000064 | -0.000061 | -0.000072 |
| 18 | 5-Oxoproline-2TMS | -0.000043 |  |  |  |  |
| 19 | Phosphoric acid-3TMS |  | -0.000197 | -0.000198 | -0.000144 |  |
| 20 | Malic acid-3TMS |  | -0.000077 | -0.000058 |  |  |
| 21 | Mannose-meto-5TMS |  | -0.000057 | -0.000055 | -0.000063 |  |
| 22 | Pyruvic acid-meto-TMS |  | -0.000065 |  |  | -0.000046 |
| 23 | Lactic acid-2TMS |  |  | 0.000062 | -0.000062 |  |
| 24 | Valine-2TMS |  |  |  |  | -0.000075 |
| 25 | Leucine-2TMS |  |  |  |  | -0.000087 |
| 26 | Isoleucine-2TMS |  |  |  |  | -0.000083 |
| 27 | Threonine-3TMS |  |  |  |  | -0.000080 |
| 28 | Glycine-3TMS |  |  |  |  | -0.000058 |
| 29 | 5-Oxoproline-2TMS |  |  |  |  | -0.000064 |
| 30 | Serine-3TMS |  |  |  |  | -0.000065 |
